# Living multicellular systems induce decodable spatial patterns in bacterial collectives

**DOI:** 10.64898/2026.09.02.748853

**Authors:** Elena G. Sergeeva, Federico Pigozzi, Thomas F. Varley, Colin J. Comerci, Zhe Zhao, Robert Brucker, Gürol M. Süel, Joshua C. Bongard, Michael Levin

**Affiliations:** Allen Discovery Center, Tufts University; Medford, MA 02155, USA; Wyss Institute for Biologically Inspired Engineering, Harvard University; Boston, MA 02115, USA; Vermont Complex Systems Institute, University of Vermont; Burlington, VT 05405, USA; Department of Computer Science, University of Vermont; Burlington, VT 05405, USA; Division of Biological Sciences, University of California San Diego; La Jolla, CA 92093, USA

**Keywords:** bacteria, *Xenopus*, embryos, Xenobots, patterns, morphogenesis, inter-kingdom, living sensors

## Abstract

Living systems continuously modify their environments through chemical, mechanical, metabolic and bioelectrical activity. Whether a presence of a multicellular system can be encoded into the emergent spatial organization of another living collective in a distributed and decodable way is unknown. Here we show that motile *Bacillus subtilis* populations reorganize their spatial and ionic collective states in response to nearby *Xenopus* embryos and Xenobots. The bacteria in a liquid culture formed autonomous motility-dependent patterns that were redirected by living targets into attraction halos, which tracked target position at a distance. Extracellular levels of potassium amplified attraction, altered local potassium dynamics, and coupled target presence to global pattern complexity. Self-supervised machine learning further identified distributed bacterial spatial signatures predictive of *Xenopus* embryo vs. Xenobot presence at a distance from the target. Together, these findings suggest that bacterial collectives can encode information about the state of other biota in their environment, revealing a previously unrecognized form of inter-kingdom interaction between living morphogenetic systems.

## 1. Introduction

Living systems continuously survey and modify their environments through chemical, metabolic, mechanical, and bioelectrical activity. These effects can extend beyond the boundaries of an individual organism, allowing the physiological state of one living system to alter the conditions experienced by another. Interactions between bacteria and multicellular organisms are therefore increasingly understood as reciprocal exchanges of information rather than as one-way microbial responses to fixed host environments [1–3]. However, it remains unclear whether information about one living system can become encoded in the emergent spatial organization of another. If so, morphology would represent not only an outcome of intrinsic self-organization, but also a distributed readout of interactions between living systems. Moreover, interactions between entities at different levels of organization (e.g., individual organisms and colonies) are especially fascinating as they bear on questions of collective decision-making and emergent dynamics at distinct scales [4–7].

Bacterial communities provide a tractable model system in which to investigate this possibility [8–12]. Although bacteria are often described as individual cells responding independently to local gradients, many bacterial populations behave as self-organizing collectives. Bacterial colonies generate reproducible macroscopic patterns through coordinated motility, growth, differentiation, secretion, metabolism, and environmental feedback [13, 14]. Biofilm formation has similarly been described as a form of microbial development involving regulated transitions among motile, adhesive, matrix-producing, and differentiated states [15]. Genetically controlled cell-state transitions and extracellular matrix components generate structured multicellular communities in which individual-cell behavior is coupled to population-scale organization [16–20]. Bacterial pattern formation can therefore be viewed as microbial morphogenesis: a process in which collective physiological state is expressed through spatial form.

Among bacterial model systems, *B. subtilis* is especially useful for examining how environmental and physiological information becomes incorporated into collective organization. In addition to transitioning between motile, sessile, matrix-producing, and differentiated states, *B. subtilis* communities coordinate behavior through electrical and ionic signaling. Biofilms use ion-channel-mediated potassium signals to synchronize metabolic states across the community [21]. These signals can alter the membrane potential and motility of distant bacterial cells, including cells belonging to other species, producing long-range attraction toward electrically active biofilms [22]. Potassium-mediated interactions can also couple separated biofilms and support temporal sharing of nutrient resources, while propagation of the signal depends on the spatial organization and participation of cells within the community [23, 24]. Bacterial communities can additionally exhibit bistable, oscillatory, and clock-like collective dynamics that organize cellular differentiation across space and time [25, 26]. These properties make *B. subtilis* collective morphology a potentially informative readout of motility, metabolic state, ionic regulation, membrane-potential dynamics, and environmental coupling.

Multicellular organisms also generate structured environments that influence microbial collective behavior. Host-derived mucus, glycans, metabolites, ions, tissue surfaces, and hydrodynamic forces can regulate bacterial adhesion, aggregation, motility, spatial organization, and biofilm formation [27–34]. Plants likewise release signals and polysaccharides that recruit beneficial bacteria and promote structured biofilm formation on roots [35–37]. Particularly relevant to the present study, epithelial potassium flux can be sensed by bacterial potassium-response machinery and promote *Pseudomonas aeruginosa* biofilm biogenesis [38]. These findings demonstrate that living tissues provide spatially and physiologically structured information that bacteria can convert into collective behavior.

Most studies in this area, however, concern direct colonization, infection, symbiosis, or bacterial organization on host-associated surfaces. Bacterial spatial structure is usually interpreted as an adaptation to a local niche and not as an information-bearing representation of the physiological state of another living system. Inter-kingdom signaling studies have shown that host hormones, microbial metabolites, immune-associated factors, and other chemical cues can mediate extensive physiological exchange between bacteria and eukaryotes [1, 2]. Microbial signals can also directly alter eukaryotic development; for example, bacterial sulfonolipids induce multicellular development in choanoflagellates [39]. The complementary possibility, that the state of a developing or engineered multicellular system may become encoded in the emergent organization of a microbial collective, has received considerably less attention.

Developmental biology provides a framework for considering how such information might enter a shared environment. Morphogenesis depends on the integration of transcriptional, biochemical, mechanical, and bioelectrical signals across multicellular networks. Endogenous ion fluxes and membrane potentials regulate proliferation, differentiation, regeneration, organ patterning in a wide range of developmental contexts [40–45]. In *Xenopus laevis*, bioelectric perturbations can regulate regenerative responses, for example, induce tail regeneration, control eye patterning, and alter tumor-like states [46–49]. Bioelectrical networks have consequently been proposed to store and process non-genetic patterning information during development and regeneration [43, 44, 50, 51]. Although these signals are generally studied for their roles within the multicellular organism, their ionic, metabolic, mechanical, and secretory consequences may also propagate into the surrounding environment and become accessible to other organisms sharing that environment.

*Xenopus* embryos and Xenobots provide related but physiologically distinct multicellular systems with which to test this hypothesis. Xenobots are synthetic living constructs assembled from embryonic *Xenopus* ectodermal cells into bodies with anatomies and behaviors distinct from those of canonical embryos [52, 53]. Their reorganization can generate coordinated locomotion and other collective behaviors, including kinematic self-replication using cells found in their environment [54]. Despite sharing a common genome and cellular origin with the embryo, Xenobots exhibit altered anatomical organization, developmental trajectory, ciliation, motility, and transcriptional state [55]. Embryos and Xenobots therefore offer a stringent comparison for asking whether bacterial collectives respond simply to the presence of *Xenopus*-derived material or can register differences in multicellular organization and physiological state of living biota in their environment.

Here, we use motile *B. subtilis*, *Xenopus laevis* embryos, and Xenobots to examine interactions between distinct living collectives embedded within a shared environment. We hypothesized that living multicellular systems alter bacterial collective organization through coupled physiological interactions, and that these interactions become expressed through changes in bacterial spatial patterning, ionic dynamics, bioelectrical state, and global pattern complexity. This framework treats bacterial populations as distributed living sensors whose emergent spatial organization may report the presence and state of nearby multicellular systems.

We used long-term fluorescence imaging of labeled *Bacillus subtilis* populations to track the emergence and reorganization of bacterial spatial patterns around *Xenopus* embryos and Xenobots. Motility-deficient and potassium-channel-deficient bacterial strains, together with passive-particle controls, were used to distinguish active microbial behavior from passive accumulation and to identify mechanisms contributing to the interaction. By varying extracellular potassium, applying physiological dyes, and comparing living targets with heat-killed, mechanically disrupted, and repositioned controls, we examined how bacterial organization responds to the physiological state, location, and ionic environment of the multicellular target. Quantitative pattern-complexity and machine-learning analyses were then used to determine whether these distributed bacterial patterns contained information about multicellular presence and biological identity.

By framing bacterial pattern formation as microbial morphogenesis, and embryos and Xenobots as interacting multicellular physiological systems, this study investigates whether the state of one living collective can leave distributed and decodable signatures within the emergent organization of another.

## 2. Results

### 2.1. Motile *B. subtilis* spontaneously generates branched spatial patterns in liquid culture

How might microbes encode information about their internal states and their environment? We first asked whether motile *B. subtilis* cells can generate macroscopic spatial organization in liquid culture in the absence of a multicellular target or other imposed spatial cue. Time-lapse imaging of *P_hyp_-yfp* motile bacteria revealed an autonomous, progressive transition from an initially dispersed fluorescence field into branched, network-like spatial patterns in the imaging plane (Fig. 1A). This behavior was not reproduced by either *Δhag P_hyp_-yfp* non-motile bacteria or passive fluorescent particles, both of which remained comparatively unstructured throughout the same analysis window.

**Figure 1.**
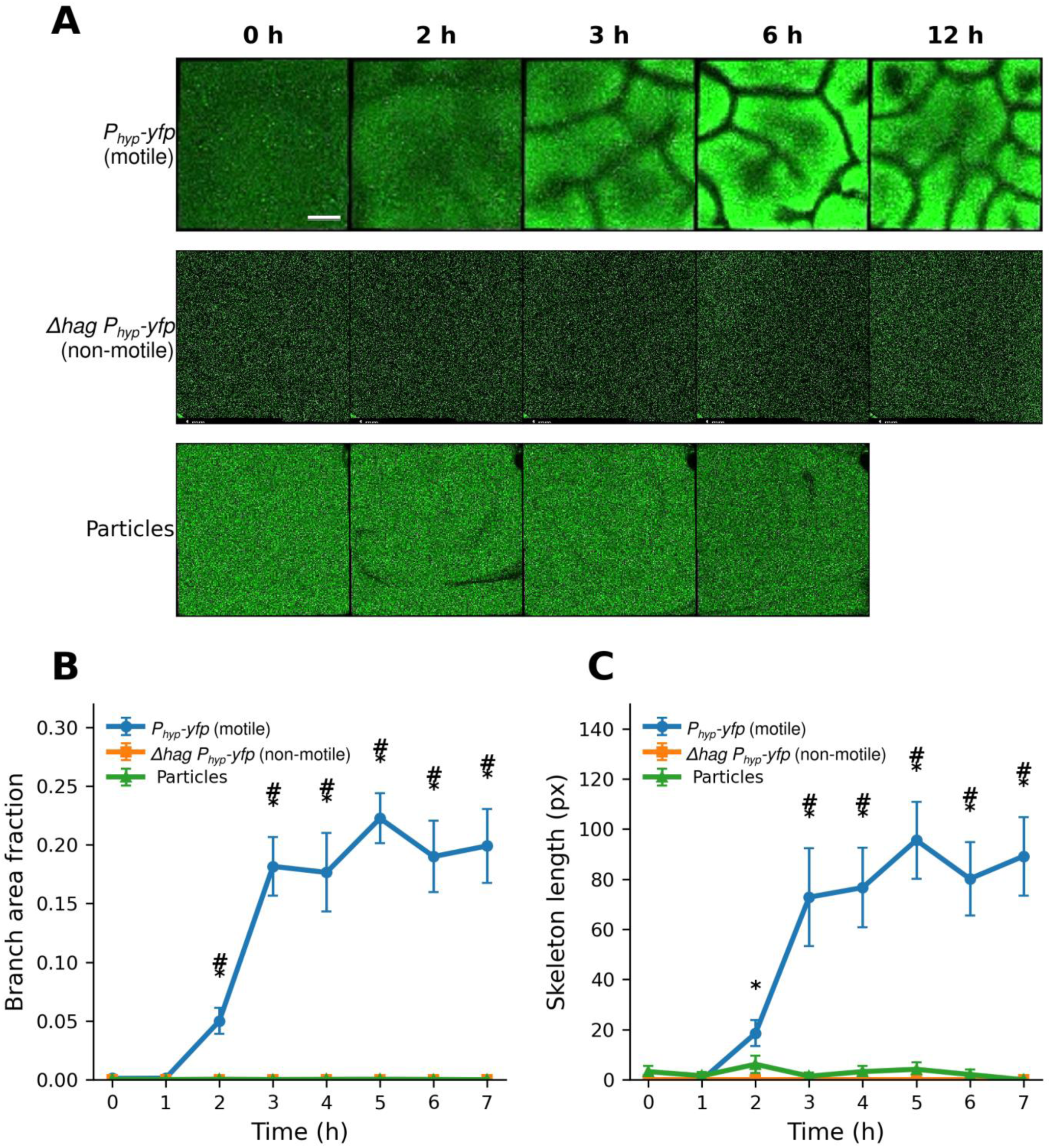
Motile *B. subtilis* cells spontaneously generate dynamic spatial patterns in liquid culture. A. Representative time-lapse fluorescence images showing pattern development in *P_hyp_-yfp* motile *B. subtilis*, *Δhag P_hyp_-yfp* non-motile *B. subtilis*, and passive fluorescent particles (1 um). Columns show the indicated time points. The motile bacterial population develops visible branched spatial organization over time, whereas non-motile bacteria and passive particles remain unbranched. Scale bar, 1 mm. B. Quantification of branch area fraction over time. Data show mean±SEM. Motile bacteria showed significant time-dependent branching dynamics over the common 0-7 h analysis window (RM ANOVA p=8.62×10⁻¹⁴; Friedman p=1.11×10⁻⁹; n=12). Non-motile bacteria showed no detectable branching at any time point (all values=0; n=8). Passive particles showed no significant temporal dynamics (RM ANOVA p=0.339; Friedman p=0.419; n=12). Area-under-the-curve (AUC) analysis over 0-7 h showed that motile bacteria differed significantly from non-motile bacteria (FDR q=1.75×10⁻⁴) and passive particles (FDR q=6.9×10⁻⁵). C. Quantification of skeleton length over time. Data show mean±SEM. Motile bacteria showed significant time- dependent increases in skeletonized branching structure (RM ANOVA p=9.97×10⁻⁹; Friedman p=3.76×10⁻⁸; n=12). Non-motile bacteria showed no detectable skeletonized branching (all values=0; n=8). Passive particles showed no significant temporal dynamics (RM ANOVA p=0.243; Friedman p=0.336; n=12). AUC analysis over 0-7 h showed that motile bacteria differed significantly from non-motile bacteria (FDR q=1.75×10⁻⁴) and passive particles (FDR q=6.9×10⁻⁵). For B-C, branch metrics were quantified from shape-filtered segmentation of elongated branch-like structures. Pairwise hour-by-hour comparisons were performed using Mann-Whitney U tests with Benjamini-Hochberg FDR correction. * indicates motile *vs Δhag P_hyp_-yfp* non-motile control, and # indicates motile vs passive particles control, FDR q<0.05.

To quantify this behavior, we measured two complementary image-based branching metrics: branch area fraction, defined as the fraction of the analyzed field occupied by segmented branch-like structures, and skeleton length, defined as the total length of the skeletonized branch network. Across the common 0-7 h analysis window, motile cultures showed a strong time-dependent increase in branch area fraction (repeated-measures ANOVA, p=8.62×10⁻¹⁴; Friedman test, p=1.11×10⁻⁹; n=12; Fig. 1B). In contrast, non-motile *Δhag P_hyp_-yfp* controls showed no detectable branch area at any time point (all values=0; n=8 wells), and passive particles showed no significant temporal dynamics (repeated-measures ANOVA, p=0.339; Friedman test, p=0.419; n=12).

Skeleton-based quantification also demonstrated a strong time-dependent increase. Motile cultures showed a significant time-dependent increase in skeleton length (repeated-measures ANOVA, p=9.97×10⁻⁹; Friedman test, p=3.76×10⁻⁸; n=12; Fig. 1C), whereas non-motile bacteria showed no detectable skeletonized branching and passive particles showed no significant change over time (repeated-measures ANOVA, p=0.243; Friedman test, p=0.336).

Direct comparison of motile bacteria with both controls further supported a motility-dependent origin of the branched patterns. Area-under-the-curve (AUC) analysis over 0-7 h showed that motile bacteria differed significantly from non-motile bacteria for both branch area fraction and skeleton length (Mann-Whitney U tests with Benjamini-Hochberg FDR correction; q=1.75×10⁻⁴ for both metrics). Motile bacteria also differed significantly from passive particles for both branch area fraction and skeleton length (q=6.9×10⁻⁵ for both metrics). Hour-by-hour comparisons showed that branch area fraction was significantly higher in motile cultures than in both controls from 2-7 h, while skeleton length was significantly higher than non-motile controls from 2- 7 h and higher than passive particles from 3-7 h.

Together, these results show that motile *B. subtilis* cells spontaneously generate dynamic, branched spatial organization in liquid culture. The absence of comparable dynamics in non-motile bacteria and passive particles shows that the observed branching is not explained by imaging background, passive particle redistribution, or static cell density alone, but requires bacterial motility. These experiments define the baseline pattern-forming capacity of motile *B. subtilis* in liquid culture. This autonomous pattern-forming behavior provides a measurable readout of bacterial collective and establishes the basis for asking what information bacterial patterns could possibly encode, for example concerning living multicellular targets in their vicinity.

### 2.2. Motile bacteria form target-associated halos around living multicellular targets

Having established that motile *B. subtilis* populations generate autonomous spatial patterns, we next asked whether these patterns could be biased by complex biological targets in the environment in a way that would allow the patterns to be a sensor for the presence of other biota.

We first tested whether bacterial collectives reorganize their spatial distribution in the presence of living multicellular targets (Xenobots). At the 1 h time point, motile *P_hyp_-yfp* bacteria formed a visible zone of fluorescence enrichment around the Xenobot, which we refer to here as a target-associated halo. This halo represented a local increase in bacterial fluorescence associated with the Xenobot boundary. Representative images showed a prominent halo in the motile *P_hyp_-yfp* condition, whereas comparable target-associated enrichment was absent in the *Δhag P_hyp_-yfp* non-motile condition and in the fluorescent particle control (Fig. 2A).

**Figure 2.**
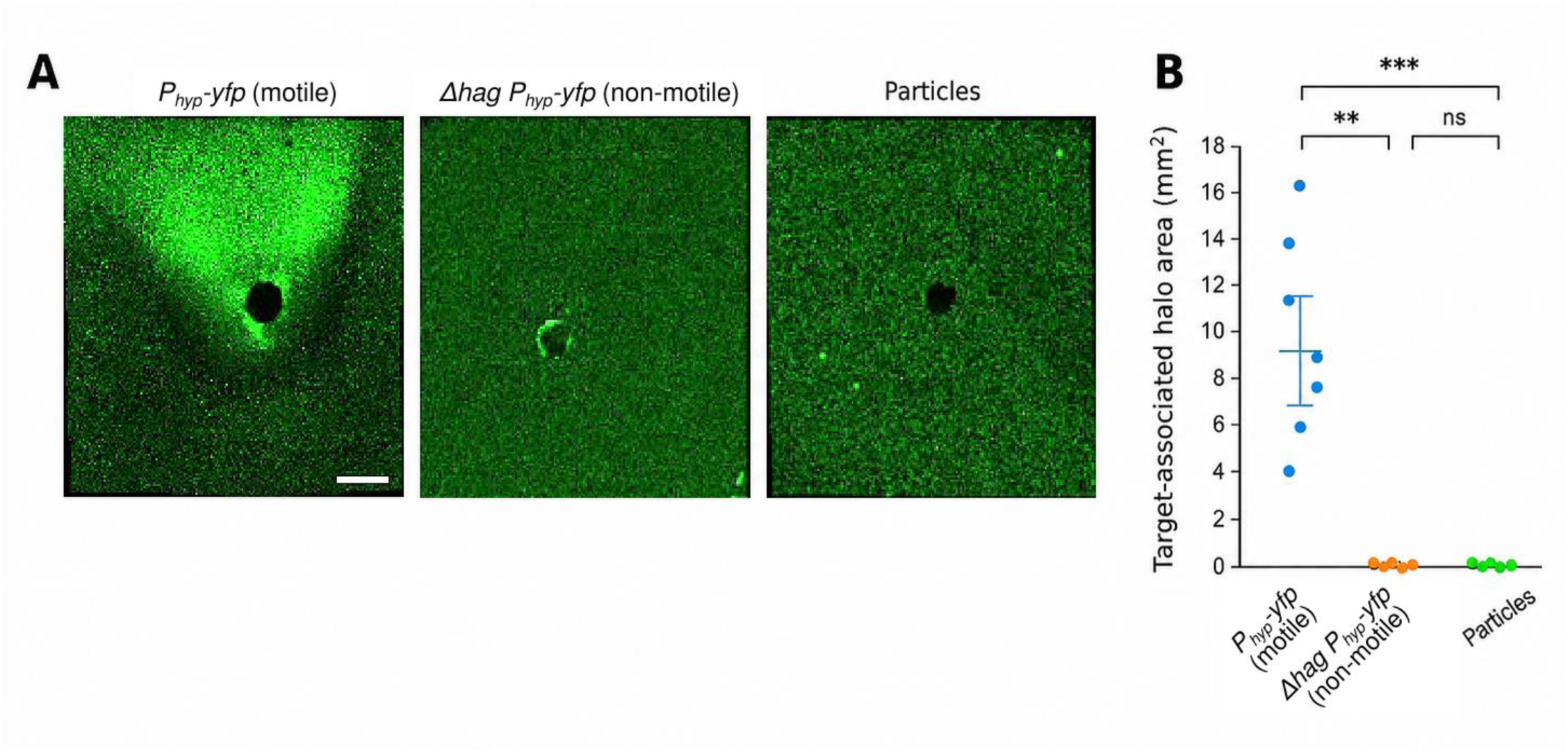
Motile *P_hyp_-yfp* bacteria form target-associated halos around Xenobots. A. Representative 1 h fluorescence images of *P_hyp_-yfp* motile bacteria, *Δhag P_hyp_-yfp* non-motile bacteria, and fluorescent particles around multicellular targets. Motile *P_hyp_-yfp* bacteria formed a prominent target- associated halo, whereas *Δhag P_hyp_-yfp* bacteria and particles showed minimal target-associated enrichment. Scale bar, 1 mm. B. Quantification of target-associated halo area in mm². Halo area was measured using a shape-independent thresholded fluorescence mask associated with the target and converted from pixels to mm² using image calibration. Each dot represents one sample (n=8 per group); bars indicate mean±SEM. Groups differed significantly overall (Kruskal-Wallis, H=15.68, p=3.93×10-4). Pairwise Mann-Whitney U tests with Holm correction showed significant differences between *P_hyp_-yfp* motile bacteria and *Δhag P_hyp_-yfp* non-motile bacteria (p=0.00185) and between *P_hyp_-yfp* motile bacteria and particles (p=4.66×10-4), but not between *Δhag P_hyp_-yfp* and particles (p=0.460). Significance labels: ns, not significant; **, p<0.01; ***, p<0.001.

To quantify this target-associated spatial reorganization, we measured halo area using a shape-independent fluorescence thresholding approach and converted the resulting area from pixels to mm². Motile *P_hyp_-yfp* bacteria produced substantially larger target-associated halos (mean±SEM: ∼9.4±1.4 mm²) than either *Δhag P_hyp_-yfp* non-motile bacteria (∼0.022±0.007 mm²) or fluorescent particles (∼0.050±0.024 mm²) (Fig. 2B). The three groups differed significantly overall (Kruskal-Wallis test, H=15.68, p=3.93×10-4). Pairwise Mann-Whitney U tests with Holm correction showed that motile *P_hyp_-yfp* differed significantly from both *Δhag P_hyp_-yfp* (p=0.00185) and particles (p=4.66×10-4), whereas *Δhag P_hyp_-yfp* and particles were not significantly different (p=0.460).

These results establish target-associated halo formation as an early bacterial collective response to multicellular targets. The absence of comparable halos in non-motile *Δhag P_hyp_-yfp* bacteria and in fluorescent particles shows that this response requires active bacterial motility and is not explained by passive accumulation of fluorescence around Xenobots.

### 2.3. Extracellular Potassium increases halo size and shifts bacterial spatial organization from branching to target- associated halo formation

Ion fluxes are often exploited by evolution for coordinating biological activity across time and space, both in microbial [21–24] and metazoan [42, 44, 45] systems. Potassium is central to *B. subtilis* physiology and bacterial bioelectrical signaling [21, 22, 64, 65]. Previous work established extracellular potassium as a mediator of long- range electrical signaling and motile-cell attraction in *B. subtilis* biofilm communities [21, 22]. We therefore asked whether extracellular potassium also modulates target-associated halo formation in our assay. To test this, we followed bacterial fluorescence around the *Xenopus laevis* embryo (NF ∼ 6-9) over an 11 h time course in KCl 1.5 mM, 100 mM, and 200 mM conditions. Representative raw time-lapse frames showed weak or minimal halo formation at KCl 1.5 mM, an intermediate halo at KCl 100 mM, and the largest halo at KCl 200 mM (Fig. 3A).

**Figure 3.**
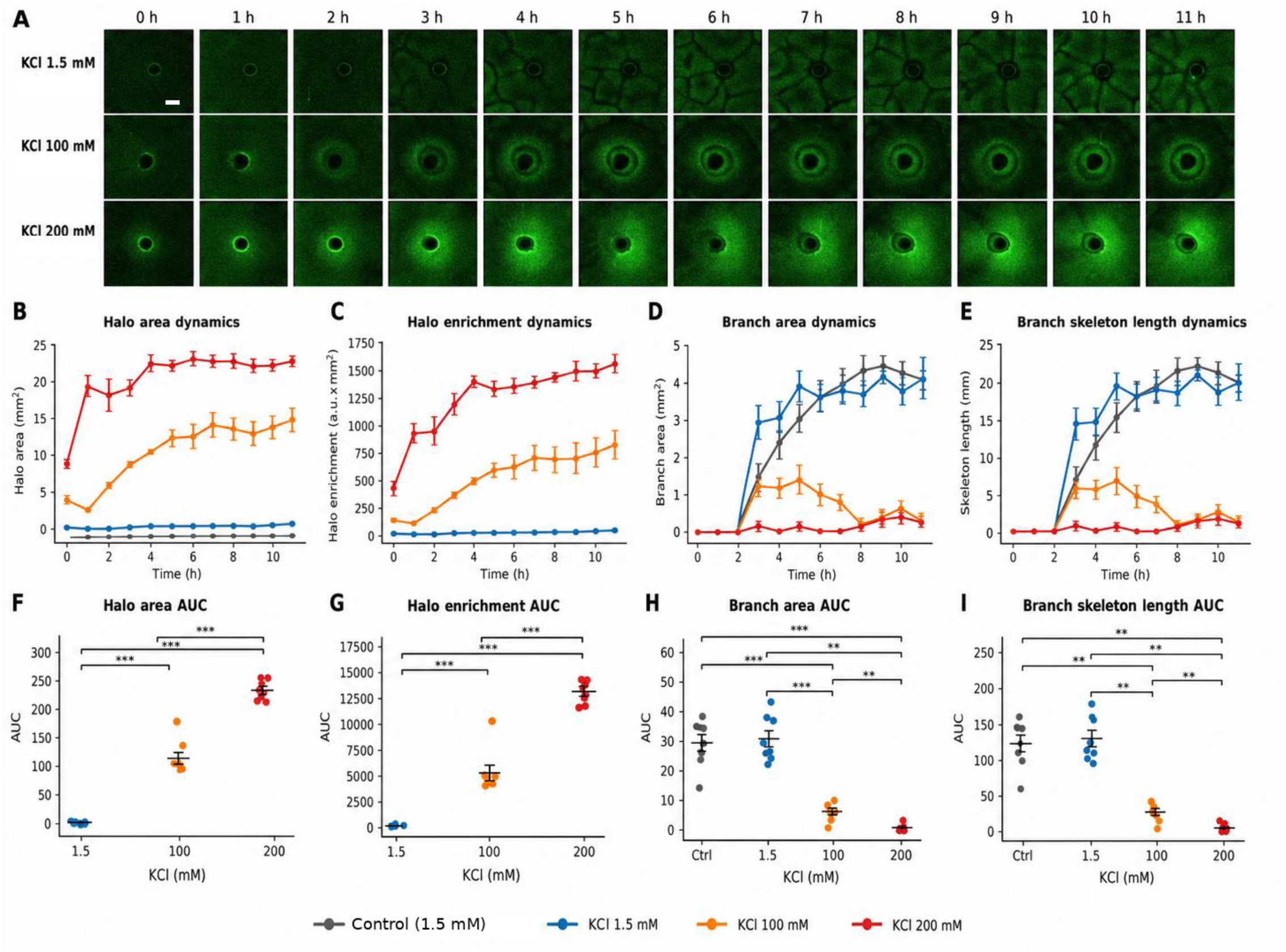
Elevated potassium enlarges target-associated halos and reduces extra-halo branching organization. A. Representative raw fluorescence frames from fixed target-containing wells followed over the 0-11 h time course in KCl 1.5 mM, 100 mM, and 200 mM. Each row shows the same well over time. Scale bar, 1 mm. B-C. Quantification of target-associated halo dynamics. Halo area (B) and halo enrichment (C) increased with KCl concentration. Halo enrichment was defined as the integrated excess green fluorescence above local background within the halo mask and is reported as arbitrary fluorescence units integrated over area, or a.u.×mm². D-E. Quantification of extra-halo branching dynamics. Branch area (D) and branch skeleton length (E) were measured outside the target-associated halo, after excluding the target and halo mask. Control denotes target-free KCl 1.5 mM wells. F-G. Area-under-the-curve analysis of halo area (F) and halo enrichment (G) across the 0-11 h time course. AUC was calculated for each well by trapezoidal integration of hourly measurements. H-I. AUC analysis of branch area (H) and branch skeleton length (I). High-KCl target-containing wells showed enlarged halos and reduced extra-halo branching compared with the target-free control and the KCl 1.5 mM target condition, consistent with a potassium-associated change in collective bacterial organization toward a target-associated halo state over a distributed branching-network state. Data are shown as mean±SEM for time-course plots and as individual wells with mean±SEM for AUC plots; n=8 wells per condition. Within-condition time effects were tested by Friedman repeated-measures tests. AUC comparisons were tested by Kruskal-Wallis tests followed by pairwise Mann-Whitney U tests with Holm correction. Significance marks: **p<0.01, ***p<0.001.

Quantification confirmed this concentration-dependent increase. Halo area increased over time and scaled with KCl concentration (Fig. 3B). At 0 h, mean halo area was 0.18±0.14 mm² at KCl 1.5 mM, 3.98±0.64 mm² at KCl 100 mM, and 9.05±0.66 mm² at KCl 200 mM. By 11 h, halo area increased to 0.67±0.20 mm², 14.93±1.61 mm², and 23.00±0.53 mm², respectively.

To capture not only halo size but also fluorescence intensity, we measured halo enrichment, defined as the integrated excess green fluorescence above local background within the halo mask. This metric is reported as arbitrary fluorescence intensity integrated over area, or a.u.×mm². Halo enrichment showed the same potassium-dependent pattern (Fig. 3C). At 11 h, halo enrichment was 32.72±13.29 a.u.×mm² at KCl 1.5 mM, 752.94±122.37 a.u.×mm² at KCl 100 mM, and 1480.44±69.70 a.u.×mm² at KCl 200 mM. Within-condition time effects were significant for halo area at KCl 1.5 mM, 100 mM, and 200 mM (Friedman p=0.0049, p<0.0001, and p=0.0001, respectively), and for halo enrichment at all three concentrations (p=0.0029, p<0.0001, and p<0.0001, respectively).

To compare cumulative halo formation across the full time course, we calculated area under the curve (AUC) for each well by trapezoidal integration of the hourly measurements from 0 to 11 h. Thus, halo area AUC represents cumulative halo area over time and is reported in mm²·h, whereas halo enrichment AUC represents cumulative excess fluorescence over time and is reported in a.u.×mm²·h. Halo area AUC increased from 3.60±0.85 mm²·h at KCl 1.5 mM to 115.46±9.75 mm²·h at KCl 100 mM and 230.79±5.39 mm²·h at KCl 200 mM (Fig. 3F). Halo enrichment AUC increased from 136.55±34.79 a.u.×mm²·h at KCl 1.5 mM to 5206.66±715.30 a.u.×mm²·h at KCl 100 mM and 13059.78±360.26 a.u.×mm²·h at KCl 200 mM (Fig. 3G). The overall KCl effect was significant for both halo area AUC and halo enrichment AUC (Kruskal-Wallis p<0.0001 for both), and all pairwise KCl comparisons were significant after Holm correction.

We next examined how potassium-enhanced halo formation related to bacterial organization outside the halo. Branching was quantified in the extra-halo compartment, defined as the analyzable well area after excluding the target and the halo mask. Branch area and skeleton length increased in the bacteria-only KCl 1.5 mM control and in the KCl 1.5 mM target condition, but remained low in target-containing wells with larger potassium-induced halos (Fig. 3D,E). In the bacteria-only KCl 1.5 mM control condition, branch area increased to 4.07±0.27 mm² by 11 h, and branch skeleton length reached 17.03±1.09 mm. The KCl 1.5 mM target condition showed a similar extra-halo branching pattern, with branch area of 4.12±0.47 mm² and skeleton length of 17.08±1.97 mm at 11 h. In contrast, at 11 h, branch area was only 0.24±0.17 mm² at KCl 100 mM and 0.20±0.13 mm² at KCl 200 mM, while skeleton length was 0.98±0.69 mm and 0.86±0.57 mm, respectively.

AUC analysis also demonstrated potassium enhanced halo formation (Fig. 3H,I). Branch area AUC was similar between the branch-only control and the KCl 1.5 mM target condition: 29.53±2.77 versus 30.89±2.63 mm²·h, with no significant difference between them (Holm-adjusted p=0.878; Fig. 3H). Branch skeleton length AUC was also similar between these two conditions: 124.39±11.65 versus 130.86±10.69 mm·h (Holm-adjusted p=0.798; Fig. 3I). In contrast, branch area AUC was lower in KCl 100 mM and KCl 200 mM target-containing wells, reaching 6.29±1.06 and 1.06±0.47 mm²·h, respectively (Fig. 3H). Skeleton length AUC was similarly reduced to 26.31±4.47 and 4.39±1.98 mm·h (Fig. 3I). The overall four-condition effect was significant for both branch area AUC and skeleton length AUC (Kruskal-Wallis p=0.000010 and p=0.000011, respectively).

To determine whether potassium itself directly altered branching dynamics, we also analyzed target-free branch-only wells across a broader KCl concentration range: 1.5, 100, 200, 300, and 400 mM (Supplement Fig. 1). Branching developed over time in these wells, but branching metrics did not show robust concentration-dependent pairwise differences after Holm correction. Branch area AUC showed only a borderline overall effect across KCl concentrations (Kruskal-Wallis p=0.0456), while skeleton length AUC and skeleton density AUC did not reach significance (p=0.0615 and p=0.0538, respectively). All pairwise KCl comparisons were non-significant after Holm correction.

Elevated potassium enlarged target-associated halos but did not globally suppress branching in target-free wells. Instead, in target-containing wells, elevated potassium is associated with a change in collective spatial organization: bacteria preferentially organize into a target-associated halo state over a distributed branching- network state.

### 2.4. Potassium sensitivity contributes to the initial stage of bacterial attraction

The potassium experiments suggested that ionic physiology contributes to embryo-directed attraction. We therefore asked whether bacterial potassium signaling components influence the timing or strength of halo formation. We focused on *yugO*, a potassium-associated channel implicated in bacterial bioelectrical dynamics [21], and compared halo formation by control *P_hyp_-yfp* cells and *ΔyugO P_hyp_-yfp* cells during co-culture with embryos.

Representative time-course images showed that both strains were capable of forming attraction halos around embryos over 12 hours, but their early dynamics differed (Fig. 4A). At 1 hour, control *P_hyp_-yfp* cells formed more prominent embryo-associated halos than *ΔyugO P_hyp_-yfp* cells. By 4 and 12 hours, the difference between strains was reduced, suggesting that loss of *yugO* delays early attraction without abolishing halo formation entirely.

**Figure 4.**
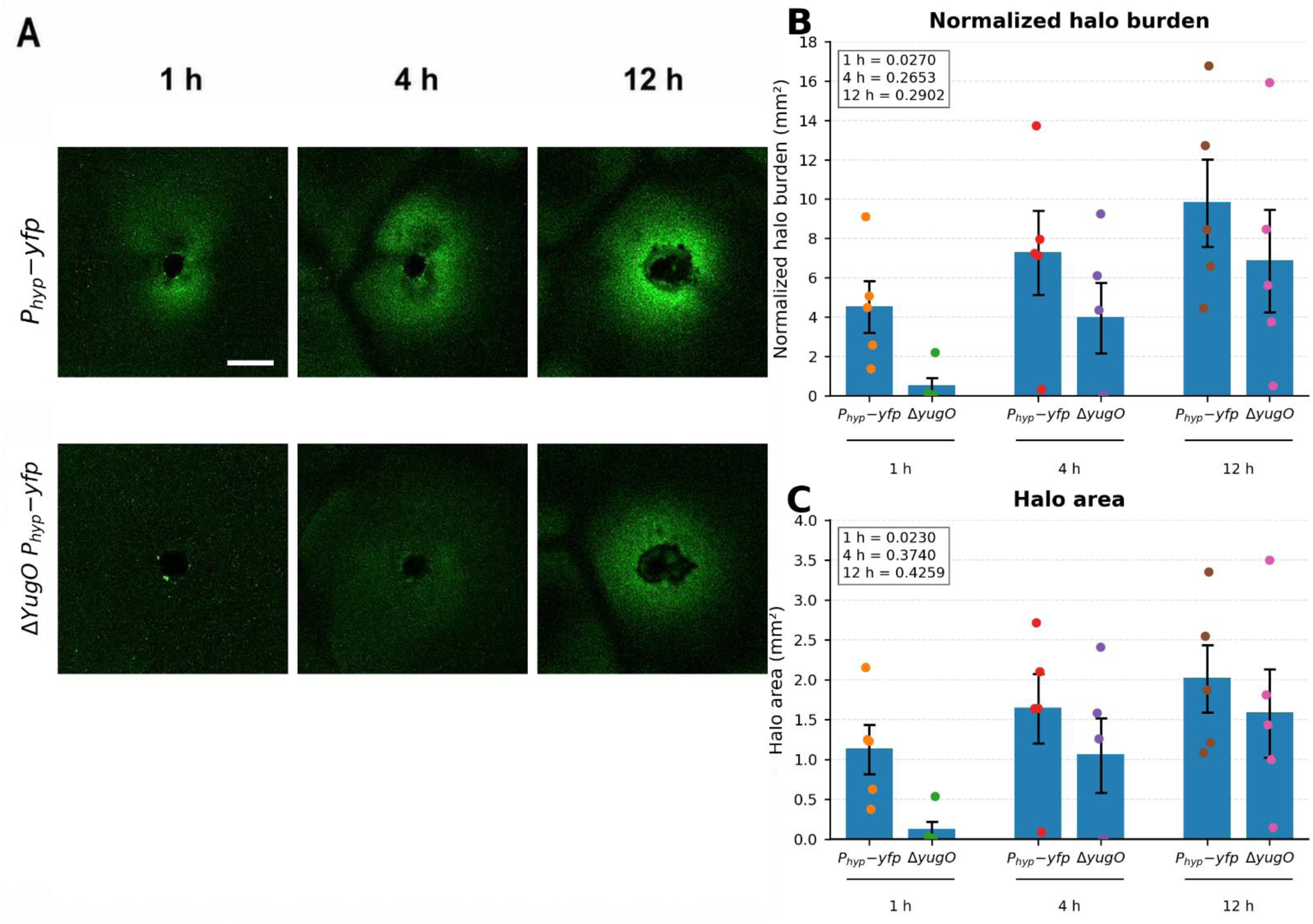
Potassium sensitivity contributes to initial stage of attraction. A. Representative fluorescence images showing bacterial halo formation around embryos at 1 h, 4 h, and 12 h of co-culture. The top row shows *P_hyp_-yfp* (WT for potassium perturbations), and the bottom row shows *Δ*y*ugO P_hyp_-yfp*. B. Quantification of normalized halo burden, calculated as halo area weighted by fluorescence enrichment relative to same-well background culture. Bars show mean±SEM, and dots show individual independent samples. Welch’s unpaired t-tests were performed separately at each selected time point. *P_hyp_-yfp* showed significantly higher normalized halo burden than *Δ*y*ugO P_hyp_-yfp* at 1 h (*p*=0.0270), while differences were not significant at 4 h (*p*=0.2653) or 12 h (*p*=0.2902). C. Quantification of halo area in mm². Bars show mean±SEM, and dots show individual independent samples. Welch’s unpaired t-tests showed a significant reduction in *Δ*y*ugO P_hyp_-yfp* halo area at 1 h (*p*=0.0230), whereas differences were not significant at 4 h (*p*=0.3740) or 12 h (*p*=0.4259). *Δ*y*ugO P_hyp_-yfp* showed delayed early halo formation, with reduced halo burden and area at 1 h followed by partial convergence with *P_hyp_-yfp* at later time points. Scale bar, 1 mm.

Normalized halo burden showed the same early difference between strains (Fig. 4B). Normalized halo burden was calculated as halo area weighted by fluorescence enrichment relative to the same-well background culture, and with embryo subtracted. At 1 hour, *P_hyp_-yfp* cells showed significantly higher normalized halo burden *than ΔyugO P_hyp_-yfp cells* (Welch’s unpaired t-test, p=0.0270). However, differences were not significant at 4 hours (p=0.2653) or 12 hours (p=0.2902).

Quantification of halo area showed a similar pattern (Fig. 4C). *ΔyugO P_hyp_-yfp* cells formed significantly smaller halos than control *P_hyp_-yfp* cells at 1 hour (p=0.0230), whereas differences were not significant at 4 hours (p=0.3740) or 12 hours (p=0.4259). Thus, the *ΔyugO* mutant exhibits a specific early-stage defect in attraction halo formation, followed by partial convergence with the control strain at later time points.

The early ΔyugO phenotype shows a contribution of potassium sensitivity to the initial stage of embryo-associated bacterial attraction. *yugO* is not strictly required for eventual halo formation, but it promotes the rapid early development of bacterial accumulation around embryos, consistent with a role for potassium-associated physiology in the onset of target-directed collective behavior.

### 2.5. Live embryonic state is required for dynamic target-associated halo formation

How much information do the bacteria have about the nature of the objects in their environment –with what specificity can the target be identified by reading their state? To test whether bacterial halo formation depends on the physiological state of the target, we compared live embryos, heat-killed embryos, and scrambled embryo material embedded in agarose under low and elevated potassium conditions. Representative raw YFP fluorescence images showed three distinct response classes (Fig. 5A). Live embryos in KCl, 200 mM produced a progressive target-associated halo that increased over the first 30 min (Fig. 5A,B). In contrast, heat-killed embryos produced a strong immediate halo-like accumulation already at the start of imaging, but this signal declined over time (Fig. 5A-C). Scrambled embryo material did not induce a compact target-associated halo under either potassium condition (Fig. 5A,B).

**Figure 5.**
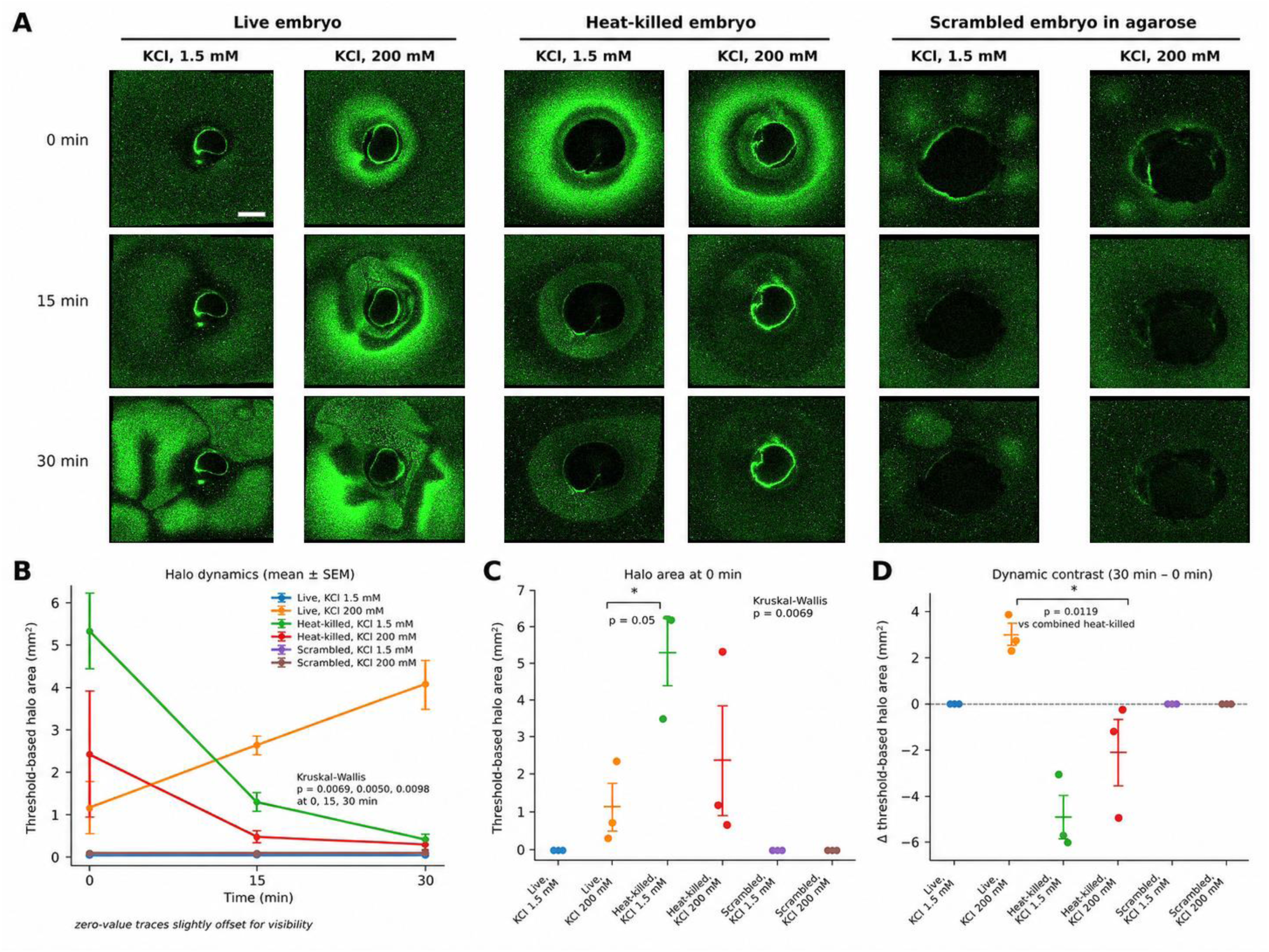
Live embryonic state is required for dynamic target-associated halo formation. A. Representative raw YFP fluorescence frames showing bacterial distribution around live embryos, heat-killed embryos, or scrambled embryo material embedded in agarose in medium containing KCl, 1.5 mM or KCl, 200 mM. Images show early time points after target exposure: 0, 15, and 30 min. Scale bar, 1 mm. B. Quantification of threshold-based target-associated halo area over time. Lines show mean±SEM, n=3 wells per condition. Zero-value traces are slightly offset for visibility only. Kruskal-Wallis tests across groups showed significant differences at 0, 15, and 30 min: p=0.0069, p=0.0050, and p=0.0098, respectively. C. Threshold-based halo area at 0 min. Heat-killed embryos in KCl, 1.5 mM showed immediate halo-like accumulation at the start of imaging, consistent with a transient accumulation response as opposed to progressive halo formation. One-sided Mann-Whitney U test versus live embryos in KCl, 200 mM, p=0.05. Kruskal-Wallis across all groups, p=0.0069. D. Dynamic contrast calculated as the change in threshold-based halo area from 0 to 30 min. Live embryos in KCl, 200 mM showed a positive change in halo area, whereas heat-killed embryos showed negative changes and scrambled embryo material remained at baseline. Dynamic, increasing target-associated halo formation was observed only with live embryos. Live embryo, KCl, 200 mM versus combined heat-killed embryo groups, one-sided Mann-Whitney U test, p=0.0119. Data are mean±SEM with individual wells shown.

Quantification of threshold-based halo area confirmed these distinct dynamics (Fig. 5B). Across the six groups, halo area differed significantly at each early time point by Kruskal-Wallis test: p=0.0069 at 0 min, p=0.0050 at 15 min, and p=0.0098 at 30 min (Fig. 4B). Live embryos in KCl, 200 mM showed an increasing halo area over time, rising from 1.15±0.63 mm² at 0 min to 2.63±0.22 mm² at 15 min and 4.07±0.59 mm² at 30 min (Fig. 5B). By contrast, heat-killed embryos showed the opposite temporal behavior: the initial halo-like accumulation was highest at 0 min and decreased over the following 30 min (Fig. 5B,C). Heat-killed embryos in KCl, 1.5 mM showed a particularly strong immediate signal at 0 min compared with live embryos in KCl, 200 mM, consistent with rapid transient accumulation as opposed to progressive halo formation (Fig. 5C; one-sided Mann-Whitney U test, p=0.05).

To distinguish progressive halo formation from immediate but transient accumulation, we calculated a dynamic contrast for each well as the change in threshold-based halo area from 0 to 30 min (Fig. 5D). Live embryos in KCl, 200 mM showed a positive dynamic contrast, whereas both heat-killed embryo groups showed negative contrasts and scrambled embryo material remained near baseline (Fig. 5D). The dynamic contrast for live embryos in KCl, 200 mM differed from the combined heat-killed embryo groups (Fig. 5D; one-sided Mann-Whitney U test, p=0.0119). Thus, elevated potassium alone was not sufficient to generate the dynamic halo response around non-living or disrupted target material. Instead, the progressive increase in target-associated bacterial organization required a live embryonic state.

### 2.6. Embryo-induced bacterial patterning dynamically tracks current embryo position

To determine whether embryo-associated bacterial enrichment reflects directional information to the embryo target, we manipulated embryo location during time-lapse imaging of *B. subtilis P_hyp_-yfp* patterning. Each well initially contained one embryo. At 2 h, embryos were either retained at the original position, supplemented with a second embryo, moved to a new position, or removed from the well (Figure 6A). Representative YFP images taken before manipulation at 1.5 h and after manipulation at 10 h showed that bacterial enrichment followed the current embryo configuration: enrichment remained around retained embryos, appeared around newly added or moved embryos, and was not maintained at sites from which embryos had departed (Figure 6B). In an exploratory maze-like multi-chamber assay, bacteria introduced into a distal chamber preferentially accumulated toward an embryo-containing well instead of a nearer empty well, consistent with longer-range embryo-associated attraction in a structured environment (Supplement Figure 2).

**Figure 6.**
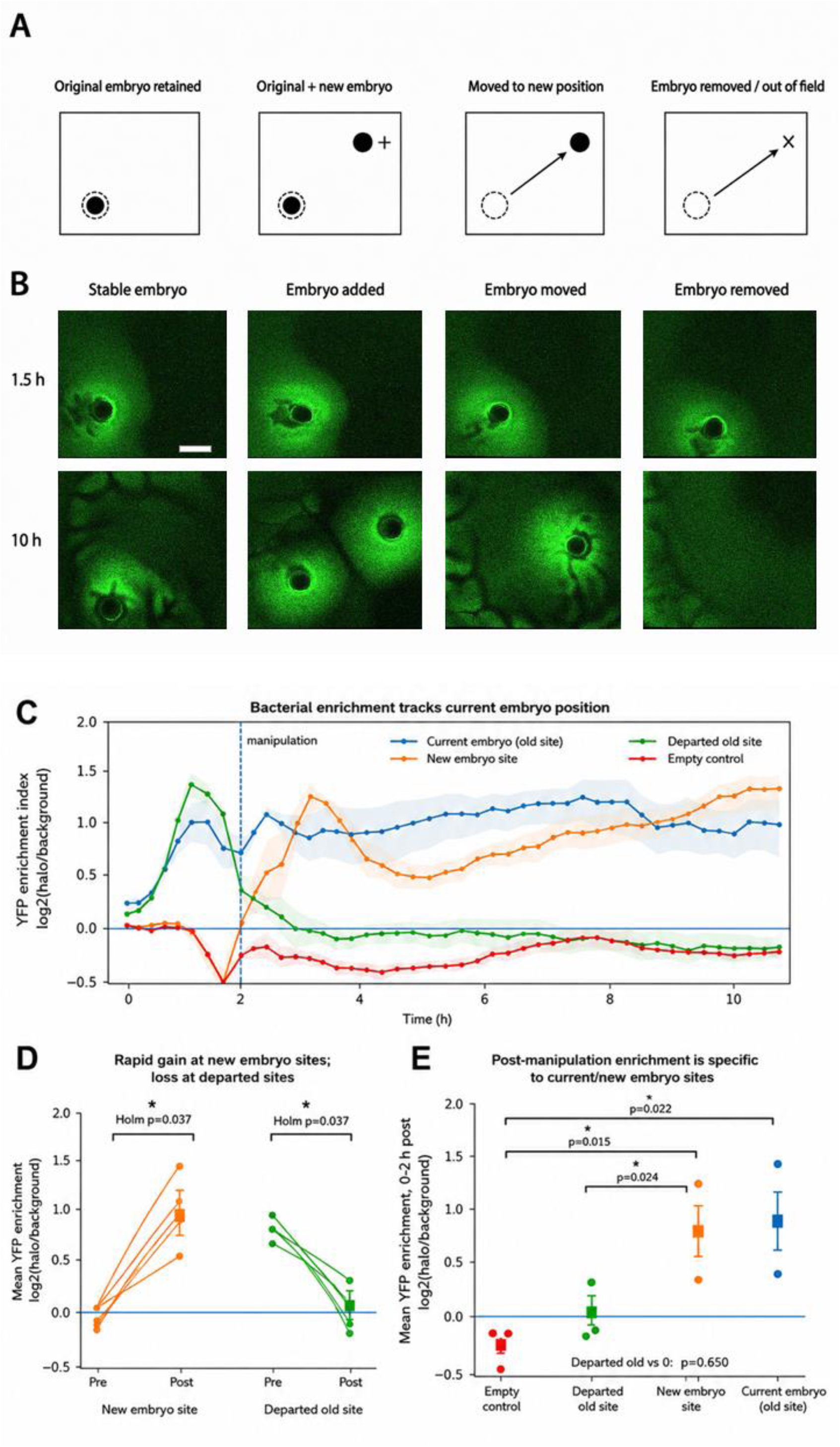
Bacterial enrichment actively tracks the current embryo target. A. Schematic of embryo-position manipulation. All wells initially contained one embryo at the original site. At 2 h, embryos were either retained at the original site, supplemented with a second embryo at a new site, moved to a new position, or removed/out of field. Dashed circles mark the original embryo position after target departure. B. Representative raw YFP fluorescence images of *B. subtilis P_hyp_-yfp* patterning before and after embryo-position manipulation. Images are shown at 1.5 h, before manipulation, and at 10 h, after manipulation. The same selected wells are shown for each condition. Scale bar, 2 mm. C. Time course of bacterial YFP enrichment at pooled site classes. Sites were grouped by target history: current embryo sites, new embryo sites, departed old sites, and empty control sites. The dashed vertical line marks embryo manipulation at 2 h. Enrichment was calculated as log2-transformed YFP intensity in the halo region relative to local background. Lines show mean±SEM. D. Paired pre/post analysis of the same physical sites before and after manipulation. New embryo sites showed increased enrichment after embryo appearance, whereas departed old sites lost enrichment after embryo movement or removal. New embryo site, post vs pre: paired t-test, Holm-adjusted p=0.037. Departed old site, post vs pre: paired t-test, Holm-adjusted p=0.037. E. Post-manipulation site-class summary during the 0-2 h interval after manipulation. New embryo sites were enriched relative to departed old sites and empty control sites, and current embryo sites were enriched relative to empty controls. New embryo site vs departed old site: Welch’s t-test, Holm-adjusted p=0.024. New embryo site vs empty control: Welch’s t-test, Holm-adjusted p=0.015. Current embryo site vs empty control: Welch’s t-test, Holm-adjusted p=0.022. Departed old sites were not significantly different from zero enrichment, one-sample t-test, p=0.650. n=4 pooled sites per class.

To quantify these dynamics, we pooled sites according to spatial history: current embryo sites, new embryo sites, departed old sites, and empty control sites. Time-course analysis showed that bacterial enrichment initially developed at original embryo positions before the 2 h manipulation. After manipulation, enrichment at departed old sites rapidly declined, whereas enrichment at new embryo sites increased and remained elevated over the subsequent hours (Figure 6C). Thus, the bacterial pattern was not fixed to a previous embryo location but instead redistributed toward the currently available embryo target.

Paired pre/post analysis confirmed this redistribution. New embryo sites increased from a mean enrichment index of −0.078 before embryo appearance to 0.824 after embryo appearance, corresponding to a mean gain of 0.902 log2 units (paired t-test, Holm-adjusted p=0.037; Figure 6D). Conversely, departed old sites decreased from 0.726 before embryo departure to 0.052 after departure, corresponding to a mean loss of 0.673 log2 units (paired t-test, Holm-adjusted p=0.037; Figure 6D).

Post-manipulation site-class comparisons provided additional evidence for active target tracking. New embryo sites were more enriched than departed old sites (Welch’s t-test, Holm-adjusted p=0.024) and empty control sites (Welch’s t-test, Holm-adjusted p=0.015; Figure 6E). Current embryo sites were also enriched relative to empty controls (Welch’s t-test, Holm-adjusted p=0.022). In contrast, departed old sites were not detectably enriched above zero after manipulation (one-sample t-test, p=0.650). Embryo-induced bacterial enrichment was reversible and actively tracked the current embryo target, dissipating from sites where the target had been moved or removed.

### 2.7. Potassium-sensitive dyes detect potassium-associated dynamics at the bacteria-*Xenopus* interface

Extracellular potassium altered bacterial localization and pattern formation around *Xenopus* embryos. We therefore asked whether those conditions might be hijacking an endogenous ionic process involved in bacteria-Xenobot signaling. To test whether potassium-related signals change near embryo-associated bacterial populations, we imaged WT *B. subtilis* (no fluorescent cell reporter) in the presence of *Xenopus* embryos using two potassium-sensitive dyes: the membrane-impermeable IPG-4 TMA⁺ Salt, used to report extracellular potassium-related signal, and the membrane-permeable IPG-4 AM, used to report intracellular-accessible potassium-related signal.

Representative images showed a potassium-dependent increase in embryo-proximal signal (Figure 7A). At 1.5 mM KCl, both dyes produced relatively weak signals around the embryo. At 200 mM KCl, IPG-4 TMA⁺ Salt showed stronger extracellular potassium-associated signal near the embryo, including localized enrichment. In embryo-only controls at high KCl, IPG-4 TMA⁺ Salt also showed local signal heterogeneity near the embryo, consistent with an embryo contribution to spatially structured extracellular potassium-associated dynamics. In parallel, IPG-4 AM showed pronounced intracellular-accessible signal accumulation in the embryo-proximal bacterial region.

**Figure 7.**
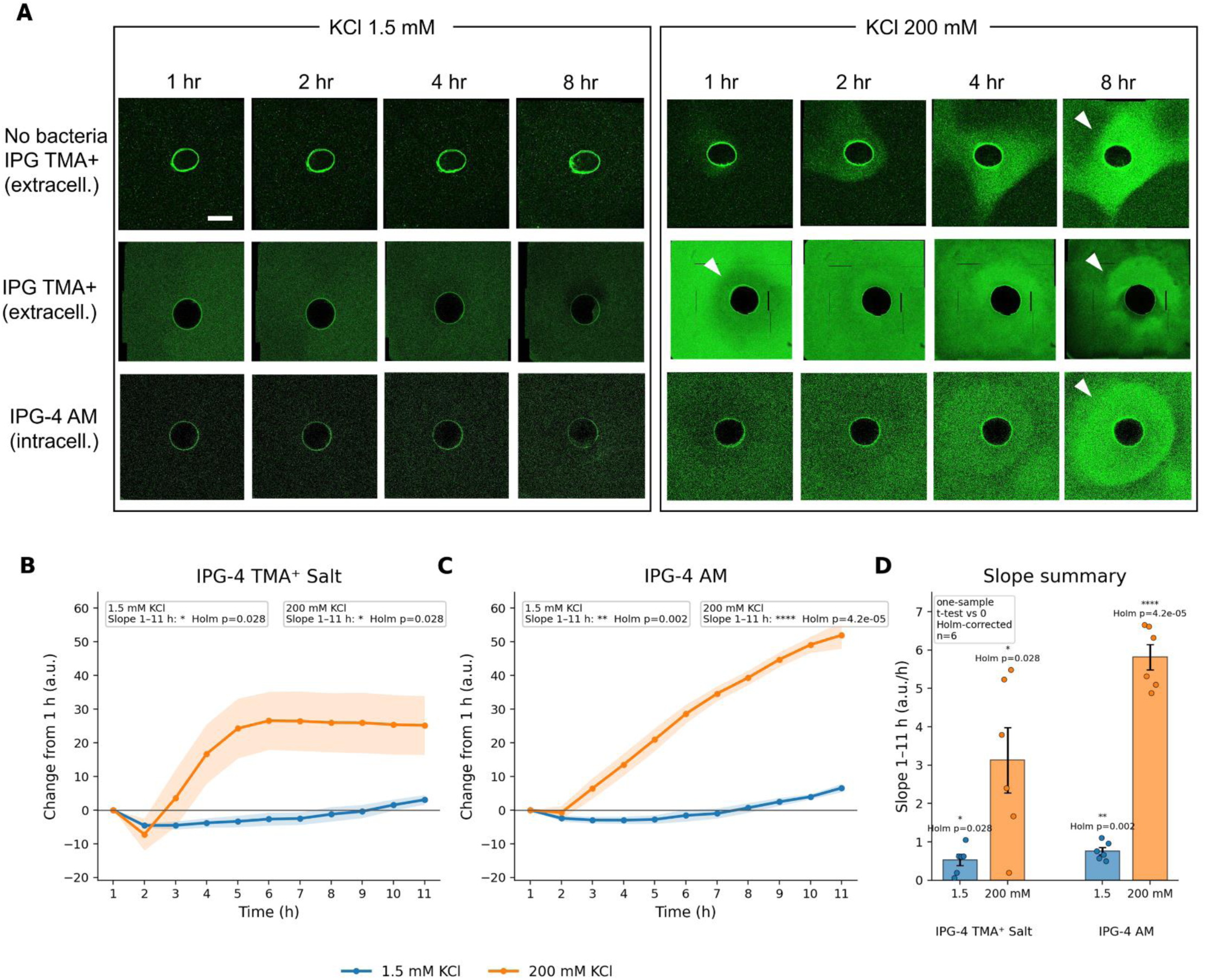
Potassium-sensitive dyes detect dynamic potassium-associated signal changes at the embryo-bacteria interface. A. Representative fluorescence images from bacteria-containing wells with WT *B. subtilis* and *Xenopus* embryos under 1.5 mM KCl or 200 mM KCl. Images show potassium-sensitive dye signal near the embryo; arrowheads mark prominent local signal enrichment near embryo-associated bacteria. Scale bar, 1 mm. B, C. Baseline-corrected near-embryo fluorescence dynamics for IPG-4 TMA⁺ Salt (B) and IPG-4 AM (C). Near- embryo signal was measured from hourly frames and expressed as change from the 1 h value. Lines show mean±SEM; n=6 repeats per condition. D. Summary of 1-11 h signal accumulation rates, calculated as the per-repeat slope of baseline-corrected near- embryo fluorescence. Bars show mean±SEM; points show individual repeats. Slopes were tested against zero using one-sample t-tests with Holm correction across the four within-condition tests. Holm-adjusted p values are shown.

To quantify these dynamics, near-embryo fluorescence was measured from hourly frames between 1 and 11 h and baseline-corrected to the 1 h signal (Figure 7B,C). To avoid frame-wise pseudoreplication, each repeat was summarized by a single planned dynamic endpoint: the slope of baseline-corrected near-embryo fluorescence from 1-11 h (Figure 7D). Each condition included n=6 repeats. Slopes were tested against zero within each condition using one-sample t-tests, with Holm correction across the four within-condition slope tests. Paired slope comparisons between KCl conditions were performed within each sensor and Holm-corrected across the two planned sensor-specific comparisons.

The extracellular sensor IPG-4 TMA⁺ Salt showed significant time-dependent signal accumulation under both potassium conditions (Figure 7B,D). At 1.5 mM KCl, the mean slope was 0.53±0.14 a.u./h (Holm-adjusted p=0.028). At 200 mM KCl, the slope increased to 3.13±0.85 a.u./h (Holm-adjusted p=0.028). Elevated KCl significantly increased the extracellular accumulation rate relative to 1.5 mM KCl (Δ slope=2.60 a.u./h; Holm-adjusted p=0.038).

The intracellular-accessible sensor IPG-4 AM showed stronger temporal accumulation, particularly under elevated potassium (Figure 7C,D). At 1.5 mM KCl, the mean slope was 0.76±0.10 a.u./h (Holm-adjusted p=0.0016). At 200 mM KCl, the slope increased to 5.81±0.33 a.u./h (Holm-adjusted p=4.2×10⁻⁵). Paired comparison confirmed that elevated KCl significantly increased the intracellular-accessible accumulation rate (Δ slope=5.05 a.u./h; Holm-adjusted p=4.0×10⁻⁵).

As a complementary analysis, we also examined whether embryo proximity was associated with bacterial membrane-potential physiology (Supplementary Figure 3). In motile *B. subtilis P_hyp_-cfp* cultures, TMRM signal transiently coincided with the embryo-associated attraction halo under high KCl. To reduce the confounding effect of motility-dependent bacterial accumulation, we also imaged non-motile Δhag *P_hyp_-yfp* cells, which do not form attraction halos. In these cells, Thioflavin and IPG-4 AM showed local membrane-potential-associated and intracellular-accessible potassium-associated signals near embryos, and time-lapse imaging showed dynamic Thioflavin signal changes over 1-8 h. These supplementary observations further link embryo proximity to local bacterial physiological dynamics, rather than bulk bacterial accumulation.

Together, these data show that embryo proximity is associated with dynamic potassium signals at the embryo-bacteria interface, and that these signals are amplified under elevated extracellular potassium. This links the potassium-dependent attraction phenotype to local ionic physiology near the embryo, supporting the idea that target-associated bacterial patterning reflects an active physiological interface and not passive accumulation.

### 2.8. Xenobot or *Xenopus* embryo presence modulates the relationship between ionic environment and bacterial pattern complexity

Showing that the bacterial population as a whole contains information about objects in their environment, we next investigated whether this information was functional for the microbial community, by testing for impacts of the *Xenopus* embryos or Xenobots on how bacteria function within their environment. In samples containing either a Xenobot or embryo, increasing KCl concentration was associated with a significant reduction in normalized LZC (r=-0.40, p<10^-4^), with bacterial spatial patterns becoming progressively less complex as KCl increased. In contrast, in samples lacking a Xenobot or embryo, no meaningful relationship was observed between KCl concentration and pattern complexity (r=0.046, n.s.), suggesting that KCl concentration alone did not systematically predict bacterial collective organization within this experimental range. This suggests that there is a three-way interaction between environment, embryo/Xenobot presence, and bacterial patterning, which can be quantified using a more sophisticated information-theoretic analysis.

When we computed the mixed datatype O-information between embryo/Xenobot presence/absence, KCl concentration, and log-transformed LZC, we found a statistically significant three-way, synergistic interaction (Ω=-0.0204 nat, p=0.0018). The negative O-information reflects a synergy-dominated higher-order dependency among the three components (two biological systems and their environment), that informs on the complexity of the patterns formed by the bacteria. Previous work has identified synergistic information with collider-like causal structures [63, 66], which in this case suggests that the bacteria are “integrating” information from both the embryo/Xenobot and their shared environment when “deciding” what kinds of patterns to create.

We next asked whether the effects on pattern complexity were subtly but detectably different in the case of Xenobots vs. unmodified embryos: do bacterial responses to environmental stressors distinguish between the presence of Xenobots vs. normal embryos? We collected a secondary dataset with fixed well-sizes and only two KCls concentrations (1.5 mM and 200 mM). Kruskal-Wallis analysis of variance found a significant difference in normalized LZC between Xenobot, embryo, and nonentity conditions (H=20.58, p<10^-4^) (Supplement figure 4). Post-hoc analysis using the Mann-Whitney U test found a statistically significant difference between the embryo and Xenobot conditions with respect to pattern complexity (U=23, p=0.00013). The embryo condition had higher LZC (0.67 +/- 0.07) than the Xenobot condition (0.55 +/- 0.046). This reveals that, not only is there an interaction between environment (KCl), bacterial organization (LZC), and entities (Xenobots and embryos), that the biological configuration of the entity in their vicinity is distinguished by the microbial community.

Together, these results show that bacterial collective organization is context-dependent. Potassium concentration, living multicellular construct status, and bacterial spatial complexity are interdependent features of the system, generating emergent collective effects detectable at the level of global spatial organization.

### 2.9. AI detected distributed bacterial pattern signatures of *Xenopus* embryo presence and distinguished between embryo and Xenobot patterns

To further characterize the latent information that the population may have about the nature of the foreign biological object in their environment, we trained a supervised vision model on fluorescence microscopy videos of bacterial cultures with three biological entities: Xenobots, embryos, or none (i.e., bacteria only). We relied on AI, due to its ability to automatically learn relevant features from data, since this is a novel model system in which relevant features are not yet known.

Since feeding the entire videos would have resulted in the AI trivially detecting the silhouette (or absence thereof) of embryos and Xenobots, we split each frame into a grid of patches and, after detecting the embryos/Xenobots, we excluded from the analysis any patch that overlapped with them. Training the AI on patches forced the model to learn a useful representation from the spatial bacterial pattern alone and ensured it could not distinguish embryo-containing cultures through silhouettes only. In other words, whatever features the AI used to analyze the bacteria’s knowledge of their environment, it would include only data from regions not including the target being sensed.

We split the 6 videos (consisting of 36 wells, across 2 KCl concentrations and 3 biological entities) between training and held-out, ensuring both splits had a balanced number of patches for each category. For statistical robustness, we trained one model for each possible split and repeated the overall procedure for 10 random seeds.

Despite the removal of direct Xenobot/embryo visual information, the classification accuracy was substantially above chance level. Accuracy on the held-out set of videos was 61.13±9.11%, significantly exceeding the performance of a 3-class majority classifier at 33% (Mann-Whitney, p<0.001). These results demonstrated that bacterial spatial organization alone contained sufficient information to predict whether a Xenobot/embryo was present in the culture. This finding showed that *Xenopus* presence induced non-local changes in bacterial collective organization that propagated throughout the surrounding microbial environment.

Moreover, we tested if the observed performance was driven by the presence of easily detectable halos right around the Xenobots/embryos. Crucially, we did not find a relationship between accuracy and distance (in number of patches) of the held-out frames from the embryo (Pearson’s r=-0.15, p=0.66), meaning that the learned features were spatially distributed and not concentrated on the thin halos. (Supplement figure 5).

Consistent with our previous results on more visible attraction patterning of KCl, we found that training on 200 mM KCl videos yielded better performance than training on 1.5 mM KCl. Indeed, average classification accuracy monotonically increased with the fraction of 200 mM KCl videos belonging to the training set (Spearman’s r=0.61, p<0.001).

Taken together, our results suggest that living multicellular systems generate measurable and spatially distributed reorganizations of bacterial pattern dynamics, producing emergent signatures detectable through representation learning even when the target systems themselves are hidden from view, and that AI tools are a tractable method for probing information content in the interactions of diverse living systems.

## 3. Discussion

We asked a fundamental question in the context of inter-kingdom interactions: how much can a bacterial community, assayed at the population level through large-scale morphogenetic readouts, detect about other living systems in its environment, and how specifically can it distinguish between them? Such readouts are useful to us as external observers because they provide an experimentally accessible means to probe bacterial sensing and discrimination and may support future applications in the emerging field of agential sensors [67, 68]. At the same time, spatial patterning is almost certainly only one subset of the information-bearing features present in the microbial population, many of which remain beyond our current ability to measure. The bacteria likely encode far more about their environment than we can presently detect. Nevertheless, even through this limited interface, we found that *Bacillus subtilis* collectives could report, at a distance, the presence, position, living state, and organizational identity of nearby *Xenopus*-derived multicellular systems. In particular, bacterial patterns distinguished frog embryos from Xenobots, two living systems derived from the same organism and genome and composed of closely related cell types, yet differing in their anatomical organization, developmental history, complexity, and internal physiological state [52, 53, 55].

The results define a progression from autonomous microbial pattern formation to target-dependent spatial organization. Motile *B. subtilis* populations spontaneously produced dynamic branched patterns in liquid culture, whereas non-motile bacteria and passive fluorescent particles did not. The patterns therefore reflected active bacterial behavior and not simple cell redistribution. In the presence of *Xenopus*-derived targets, this autonomous collective organization was redirected into target-associated halos. Halo formation required bacterial motility, increased with extracellular potassium, tracked changes in target position, and depended on the physiological state of the target. Live embryos produced progressively developing halos, while heat-killed embryos produced strong but transient early accumulation and scrambled embryonic material did not produce the same organized response. At larger spatial scales, *Xenopus* target presence altered the relationship between potassium concentration and bacterial pattern complexity, and machine learning detected distributed bacterial signatures of embryo versus Xenobot identity even after the targets themselves were excluded from the images.

These findings build on a long-standing view of bacterial populations as self-organizing, development-like systems [13–20]. Host- and biota-derived environments are already known to shape bacterial organization through more than generalized effects on growth [27–38]. Here, bacterial collective morphology is examined as a distributed and decodable readout of a nearby multicellular system. The bacteria did not simply accumulate on a tissue surface: their spatial organization tracked a target across the shared environment, responded differently to live and non-living target states, and contained information about target identity beyond the immediate target boundary. Thus, the novelty lies not simply in showing that a biological environment alters bacterial organization, but in demonstrating that bacterial organization can report multiple features of another living system, including its presence, current position, physiological activity, environmental coupling, and multicellular configuration.

The potassium dependence of this interaction extends prior work showing that *B. subtilis* biofilms use potassium-mediated bioelectrical signaling to coordinate collective physiology and long-range behavior [21–26]. Our system differs from these bacterial biofilm paradigms because the relevant multicellular target is not another bacterial community but a *Xenopus* embryo or Xenobot. Nevertheless, the convergence on potassium suggests that bacteria may use conserved ionic mechanisms to respond to physiological perturbations generated by very different forms of life. Elevated KCl increased target-associated halo formation and shifted bacterial organization away from distributed branching toward target-associated accumulation. Importantly, potassium did not robustly suppress branching in target-free controls, arguing against a simple global collapse of spatial organization. Instead, potassium changed how the bacterial collective responded to the target.

The *ΔyugO* strain, which lacks the *YugO* potassium channel implicated in *B. subtilis* bioelectrical signaling [21], further refined this interpretation. Loss of *YugO* delayed early halo formation but did not prevent eventual attraction. Thus, *YugO*-dependent physiology appears to facilitate the rapid onset or amplification of the response rather than acting as an indispensable receptor for a single target-derived signal. This partial phenotype is consistent with a distributed mechanism in which potassium signaling interacts with chemotaxis, membrane potential, metabolism, osmotic regulation, or other physiological pathways. Potassium may therefore function as an enabling or amplifying component of the interaction and not as the sole instructive cue. This interpretation is consistent with more complex organization of potassium homeostasis in *B. subtilis*, which involves multiple uptake and regulatory systems [64, 65].

The potassium-sensitive dye measurements provide a physiological bridge between the potassium-dependent behavioral phenotype and local conditions at the embryo-bacteria interface. Both extracellular and intracellular-accessible potassium-associated signals increased near embryos, and their temporal accumulation was enhanced under elevated extracellular potassium. Membrane-potential-associated changes were also observed near embryos, including in non-motile bacteria that did not form attraction halos. These observations suggest that embryo proximity is associated with local ionic and bioelectrical dynamics that cannot be explained solely by the bulk accumulation of motile cells. These observations suggest that living *Xenopus*-derived targets reshape the local physiological environment and bacteria convert this information into collective spatial organization. However, the dye measurements do not yet establish the direction of potassium flux, identify its cellular source, or demonstrate that potassium alone is sufficient to specify the observed patterns.

This interpretation is especially relevant in the context of developmental bioelectricity. Endogenous membrane potentials and ion fluxes regulate proliferation, differentiation, regeneration, organ patterning, and tumor state across multicellular systems [40, 41, 43, 44, 46–51]. Our findings raise the possibility that aspects of these physiological states are not only used internally by a multicellular system, but can also become externally legible to another living collective. This does not yet demonstrate that bacteria directly decode a specific embryonic bioelectric pattern. Rather, it suggests that the ionic and physiological outputs of morphogenesis can propagate into the shared environment and become transformed into bacterial spatial form.

The live-target and tracking experiments strengthen this interpretation. Progressive halo formation required a living embryo, while heat-killed embryos produced a qualitatively different, transient response. This distinction argues against a response driven solely by tissue composition, fixed geometry, or *Xenopus*-derived biomass. Moreover, when embryos were moved, added, or removed, bacterial enrichment reorganized toward the current target configuration and dissipated from departed locations. The spatial pattern therefore remained dynamically coupled to present conditions instead of preserving only a static history of earlier target positions. The response was therefore active, reversible, and dynamically coupled to target-associated physiology.

The embryo-Xenobot comparison tested whether the bacterial collective can distinguish between closely related multicellular systems. Xenobots are reconfigurable living systems assembled from *Xenopus* embryonic cells into bodies with forms and behaviors that differ from those of canonical embryos [52, 53]. Such constructs can exhibit coordinated locomotion and even kinematic self-replication through collective manipulation of dissociated cells [54]. Their altered organization and life history are also accompanied by transcriptional differences despite the absence of genomic engineering [55]. Embryos and Xenobots therefore share biological material and genome while differing in anatomical architecture, developmental trajectory, ciliation, motility, and physiological organization.

These differences were reflected in bacterial patterning. Embryo- and Xenobot-containing conditions produced distinguishable effects on bacterial pattern complexity, and the machine-learning classifier discriminated embryo, Xenobot, and bacteria-only conditions above chance after direct visual information from the targets had been removed. This suggests that the bacterial response is not limited to a generic detection of *Xenopus* tissue. Instead, features of multicellular organization appear to propagate into the surrounding bacterial field.

This observation also connects to a broader framework, where behavior in embodied systems emerges from the coupling of morphology, control, and environment, with no single controller acting in isolation. [69–72].

Xenobots demonstrate this principle in living material: changing the organization of cells changes the behavioral and functional possibilities of the resulting body [52–54]. Our findings add a complementary dimension. The organization of a living body may affect not only its own behavior, but also how it is registered by other living systems. A bacterial collective can therefore function as a distributed observer of embodiment, converting otherwise subtle differences in multicellular organization into measurable changes in population-scale form.

The pattern-complexity analysis further indicates that this interaction cannot be reduced to a simple effect of potassium or *Xenopus* target presence alone. Increasing KCl was associated with reduced bacterial pattern complexity when an embryo or Xenobot was present, but not in bacteria-only conditions. The significant negative O-information reflects a synergy-dominated higher-order dependency among target presence, potassium concentration, and bacterial complexity [63, 66]. This does not by itself establish a causal architecture, but it shows that information about the system is generated jointly by the target, bacterial population, and shared ionic environment. The resulting pattern therefore belongs to the coupled system and not to any one component considered in isolation.

The machine-learning analysis reached a related conclusion from a different direction. Even after embryo- and Xenobot-containing regions were excluded, bacterial-pattern patches retained sufficient information for above-chance classification. The lack of a detectable relationship between classification accuracy and distance from the target further suggests that the informative features were not restricted to the visible attraction halo. Instead, target-associated state was represented across a broader portion of the bacterial field. This connects the study to image-based phenotypic profiling, in which high-dimensional biological states can be recovered from spatial features that are not readily captured by predefined metrics or human inspection [73, 74]. Here, artificial intelligence is not evidence that the mechanism has been understood; it is evidence that distributed biological information exists in the bacterial morphology and can be decoded.

### Limitations

Several technological limitations leave important questions for subsequent work. First, relevant dynamics in ecological settings remain to be measured; the elevated KCl concentrations used here are experimental perturbations and should not automatically be interpreted as normal physiological conditions. Second, we do not claim that the dynamics we characterized are the entire mechanism or signature of interaction – it is very likely that many more dimensions of physiology and biophysical events remain to be discovered. Third, potassium- and voltage-sensitive dyes provide relative physiological readouts rather than direct measurements of ion flux or membrane voltage, and their signals may be affected by cell density, dye loading, optical geometry, and background fluorescence. Improved sensors, covering a broader set of physiological events, will surely provide better future insight into the full richness of inter-kingdom communications at many scales. Finally, the pattern-complexity and O-information analyses identify statistical structure but do not specify mechanism or causal direction. In the machine-learning study, feature-attribution and perturbation approaches will be needed to identify which aspects of bacterial organization support classification and whether those features are mechanistically involved in the biological response.

### Future directions

Expanded panels of bacterial ion-channel, chemotaxis, motility, and metabolic mutants could identify the sensing and response pathways required on the microbial side. Early transcriptomic and metabolomic measurements could reveal physiological changes preceding visible pattern formation. Future work should include both, a delineation of the molecular biology of signaling transduction cascade, and a semantic analysis akin to neural decoding [75, 76] to translate exactly what these living beings are saying to each other. Microfluidics and closed-loop control systems could learn to exploit these dynamics to make predictions in useful ecological or biomedical settings. What other states of living tissue could bacterial colonies inform us about, as agential sensors? Comparing embryos of different stages, healthy and perturbed embryos, distinct Xenobot designs, and multicellular tissues of other species would establish which properties of living systems are represented in bacterial collective form.

### Conclusion

Our results suggest a broader way to think about interactions among living systems. A multicellular body does not only construct and maintain its own morphology. Through its ionic, metabolic, mechanical, secretory, and bioelectric activity, it also modifies the state space available to nearby organisms [77–79]. A bacterial population, in turn, does not merely respond as a collection of independent cells, but integrates these environmental changes into collective spatial organization. Morphogenesis can therefore serve as an interface between highly diverse systems: the physiological state of one collective becomes partially written into the form of another. This perspective extends host-microbe interaction beyond colonization and molecular exchange toward distributed, decodable coupling between morphogenetic systems, and suggests a foundation for living sensors capable of reporting biological states that remain difficult to detect by conventional means.

## 4. Materials and Methods

### 4.1. Experimental design

The study examined spatial and physiological responses of *Bacillus subtilis* populations to *Xenopus laevis* embryos and Xenobots. The experimental series included target-free bacterial cultures (without embryos or Xenobots), motility-deficient bacterial controls, passive fluorescent-particle controls, extracellular and physiological potassium manipulations, live and non-living target preparations, embryo-position manipulations, confocal time-lapse imaging of fluorescent cell reporters, extracellular and intracellular potassium sensors, and membrane-potential-sensitive dyes, bacterial pattern-complexity analysis, and machine-learning classification of distributed bacterial patterns.

The independent experimental unit was an independently prepared well with bacterial liquid culture with or without target for image-based time-lapse biological experiments and a source video for computational experiments. Time frames from the same well video were treated as repeated observations. Exact sample sizes and analysis windows are specified for each experiment.

### 4.2. Biological preparations

#### Xenopus laevis embryos

Adult *Xenopus laevis* were maintained at Tufts University, Levin Lab frog facilities, and all animal procedures were approved by the Tufts University Institutional Animal Care and Use Committee under protocols M2023-18 and M2026-13. Embryos were generated by standard *in vitro* fertilization, reared at 14 °C in 0.1X Marc’s Modified Ringer’s solution (MMR; pH 7.8), and staged according to Nieuwkoop and Faber criteria [56]. Embryos from separate females were pooled and randomly assigned to experimental groups at NF stages 9-14 unless otherwise indicated.

#### Xenobots

Xenobots were generated from animal-cap ectoderm dissected from NF stage 9 *Xenopus* embryos using established procedures [52]. Stage 9 embryos were transferred to Petri dishes coated with 1% agarose prepared in 0.75X MMR and containing 0.75X MMR. After removal of the vitelline membrane, animal-cap tissue was dissected and placed on the agarose surface with the inner face upward. Over approximately 2 h, the explants rounded into spherical tissue constructs. Constructs were then cultured at 14 °C on 1% agarose in 0.75X MMR with daily medium changes, allowing maturation. Depending on the experiment, either immobile unciliated

Xenobots (Day 2-3 after generation, before cilia differentiation) or motile ciliated Xenobots (Day 5-6 after generation) were used in bacterial interaction assays.

#### Bacterial strains

All experiments were performed using *Bacillus subtilis* NCIB 3610 derived strains [57, 58]. Most experiments used a motile *P_hyp_-yfp* reporter strain, in which YFP was expressed under the *Phyperspank* promoter to label the bacterial population. Non-motile *Δhag P_hyp_-yfp* bacteria were used in specified control experiments to test the requirement for flagellar motility. To probe physiological potassium, *ΔYugO P_hyp_-yfp* strain with deleted *YugO* potassium ion channel was used. For potassium-sensitive dye experiments, wild-type NCIB 3610 bacteria were used to avoid spectral overlap with the YFP reporter. For TMRM imaging, a *P_hyp_-cfp* reporter strain was used instead, enabling simultaneous visualization of bacterial distribution without interference from YFP fluorescence.

#### Bacterial culture and assay medium

Cells were streaked from −80 °C glycerol stocks onto LB Agar, Miller (Fisher BioReagents, BP1425) and incubated overnight at 37 °C. A single colony was inoculated into 2 ml LB Broth, Miller (Fisher BioReagents, BP1426) containing the appropriate strain-specific antibiotics for selection (*P_hyp_-yfp*: spectinomycin, 300 µg/mL, Sigma-Aldrich, S4014; Δ*hag P_hyp_-yfp*: chloramphenicol, 5 µg/mL, Sigma-Aldrich, C0378; *ΔyugO P_hyp_-yfp*: kanamycin, 10 µg/mL, Sigma-Aldrich, K1377, together with spectinomycin 300 µg/mL) and grown at 37 °C with shaking for several hours, typically to OD600≈2.5. Cultures were centrifuged at 2100xg for 3 min, washed, centrifuged again, and resuspended to OD600=0.5 in minimal co-culture medium containing 0.1X MMR, 0.5% (w/v) monosodium L-glutamate (Sigma-Aldrich, G1626), and 0.5% (v/v) glycerol (Fisher BioReagents, BP229). Isopropyl β-D-1-thiogalactopyranoside (IPTG; Sigma-Aldrich, I6758) was added at final concentration 1 mM to induce reporter expression and for bacteria to adapt to the medium conditions. Cultures were then cooled to room temperature before addition of *Xenopus* embryos, Xenobots, dyes, particles, or other experimental components. Potassium chloride (Fisher BioReagents, BP366) was prepared as a 1M stock and added in specified concentrations immediately before imaging.

#### Non-living Xenopus embryo controls

Stage-matched embryos were heat-killed by incubation in pre-equilibrated 0.1X MMR at 48 °C for 10 min. For scrambled-tissue controls, stage-matched embryos were mechanically dissociated by repeated pipetting until no visible anatomical organization remained. The homogenized tissue suspension was mixed with low melting point agarose (to final 1.5% agarose concentration; Invitrogen, Thermo Fisher Scientific,16520) prepared in minimal co-culture medium and deposited as localized gel cubes.

### 4.3. Co-culture conditions, controls, and time-lapse imaging

Embryos or Xenobots were placed near the center of imaging wells containing bacterial cultures using a transfer pipette unless otherwise specified; target-free controls contained bacteria alone. Slides were covered with air-permeable lids during imaging. Passive controls used 1-µm fluorescent particles adjusted to produce a measurable fluorescence field. Experiments were performed in Ibidi imaging slides using 150 µl per well for µ- Slide 18 Well (Ibidi GmbH, 81816), 500 µl per well for µ-Slide 8 Well (Ibidi GmbH, 80826), and 1500 µl per well for µ-Slide 2 Well (Ibidi GmbH, 80286). Time-lapse imaging was performed on Leica Stellaris/SP8 inverted confocal microscopes with environmental control. Most experiments used a 10X objective, 1X digital zoom, a 3- Airy-unit pinhole, laser power between 2 and 10 depending on the fluorophore and experiment, bidirectional scanning at speed 400, and sequential channel acquisition. For cell resolution imaging, 20X objective, 4X digital zoom, a 1-Airy-unit pinhole settings was used. Bright-field images were acquired in parallel with fluorescence images. Tiling and stitching were used to image entire wells. Nominal excitation/emission settings were 515/530 nm for YFP, 433/475 nm for CFP, 525/545 nm for the potassium-sensitive dyes IPG TMA+ and IPG-4 AM, 450/485 nm for Thioflavin T, and 550/575 nm for TMRM. Samples were maintained at 18 °C during imaging. Image acquisition time-lapse schedules were experiment specific and noted for each experiment. Pixel areas and distances were converted to physical units using image-specific calibration.

### 4.4. Branch-pattern segmentation and quantification

Branching was quantified from the green fluorescence channel using a robust, multistage curvilinear- feature segmentation pipeline (see 4.10 for the details). Each frame was intensity-normalized and corrected for slowly varying background by subtracting a large-scale Gaussian-smoothed background. Branch-like features were enhanced using multiscale morphological black-hat filtering, contrast-limited adaptive histogram equalization, and a Frangi ridge response. Candidate branch pixels were then selected by adaptive hysteresis thresholding. The binary mask was skeletonized, short isolated fragments were removed, and connected components were retained only when their skeleton topology was network-like. Thus, component selection was based on network structure instead of a single fixed aspect-ratio, solidity, or width cutoff. Images were segmented without cropping or resizing, preserving the original field of view.

Branch area fraction was calculated as the number of pixels in the retained branch mask divided by the number of analyzable pixels. For calibrated datasets, branch area was reported in mm². The retained mask was skeletonized to a one-pixel-wide representation, and total skeleton length was calculated from the skeletonized network and measured in pixels or converted to millimeters. Skeleton density was calculated as total skeleton length divided by branch area.

For each well, area under the curve (AUC) was calculated over the specified time window by trapezoidal integration. Branch-area AUC was expressed in mm²·h and skeleton-length AUC in mm·h using physical calibration.

### 4.5. Target-associated halo analysis

A target-associated halo was operationally defined as a shape-independent region of elevated bacterial fluorescence spatially associated with the target’s (embryo, Xenobot or control target) boundary. The target body was omitted from the analysis, and the halo mask was not constrained to a circular geometry. Halo area was calculated as the threshold-positive area outside the exclusion region and converted from pixels to mm² using the calibration for the corresponding image series (see 2.10 for pipeline details). Mean halo enrichment was calculated as the mean fluorescence within the halo mask divided by the mean same-well culture fluorescence, and normalized halo burden was calculated as halo area×mean halo enrichment. When no halo was detected, halo area and normalized halo burden were assigned a value of zero. In analyses of integrated excess halo signal, fluorescence above the same-well background was integrated over the calibrated halo area and reported in arbitrary fluorescence units×mm². For extra-halo branching analysis, the target and halo masks were removed before branch area and skeleton length were quantified. For each well, area under the curve was calculated by trapezoidal integration over the specified time window; halo-area AUC was expressed in mm²·h, and integrated excess halo-signal AUC in a.u.×mm²·h.

### 4.6. Embryo position manipulation and spatial tracking

Embryo-position manipulation assays were performed in larger 8-well Ibidi slides using minimal co-culture medium containing 200 mM KCl. Each well initially contained one embryo positioned off-center, usually near the lower-left region of the well. At 2 h, embryos were either retained at the original position, supplemented with a second embryo at a new position (upper-right region of the well), moved to a new position with a pipette tip, or removed using a cut 1-ml pipette tip. Sites were classified by target history as current embryo, new embryo, departed old, or empty control sites. For each site and time point, bacterial enrichment was calculated as the log2-transformed ratio of YFP intensity in the site-associated halo region to local background intensity. The same physical sites were retained for paired before-versus-after analyses.

Long-range attraction was examined in custom maze-like multi-chamber geometries made by modifying Ibidi imaging dishes containing four-chamber silicone inserts (Ibidi, 80409). Approximately 0.5-mm-wide openings were cut between chambers to create liquid-connected paths. In the four-chamber configuration shown in Supplementary Figure 2, an embryo was positioned in the upper-left distal chamber and *P_hyp_-yfp* bacteria were introduced into the bottom-right chamber; bacteria could redistribute through the connected chamber network toward the target-containing or empty chambers. Fluorescence was monitored for up to 12 h. The displayed assay was interpreted descriptively and was not included in inferential testing.

### 4.7. Potassium-sensitive and membrane-potential-associated imaging

Potassium-sensitive imaging used IPG-4 TMA+ Salt (ION Biosciences, 3023F) as a membrane-impermeant reporter of extracellular potassium-associated signal and IPG-4 AM (ION Biosciences, 3021F) as a membrane-permeant reporter of intracellular-accessible potassium-associated signal. Wild-type NCIB 3610 *B. subtilis* lacking a fluorescent cell reporter was imaged with *Xenopus* embryos in minimal co-culture medium containing 1.5 or 200 mM KCl; embryo-only controls were included for the extracellular reporter. Dyes were prepared according to the manufacturer’s instructions as DMSO stock solutions and added immediately before imaging to a final concentration of 3 µM. Samples were imaged using nominal 525/545 nm excitation/emission settings.

For each movie, mean fluorescence was extracted from an embryo-proximal region while excluding the dark embryo body. Hourly values from 1 to 11 h were baseline-corrected to the 1-h value, ΔF(t)=F(t)-F(1h). Each experimental repeat was reduced to one dynamic endpoint by fitting an ordinary least-squares line to the baseline-corrected 1-11 h values; the fitted slope in a.u./h was used for statistical analysis. Six repeats were analyzed per sensor and KCl condition. KCl conditions were paired within each sensor.

Supplementary membrane-potential-associated imaging used tetramethylrhodamine methyl ester (TMRM; Invitrogen, Thermo Fisher Scientific, T668) with motile *P_hyp_-cfp* bacteria in 300 mM KCl and Thioflavin T (ThT; Acros Organics, AC211761000) with non-motile Δ*hag P_hyp_-yfp* bacteria in 1.5 mM KCl. IPG-4 AM was also imaged in the non-motile condition. TMRM, ThT, and IPG-4 AM were prepared as DMSO stocks and added to the samples immediately before imaging. TMRM was used at a final concentration of 200 nM and imaged using nominal 550/575 nm excitation/emission settings, while CFP fluorescence was acquired using nominal 433/475 nm excitation/emission settings. ThT was used at a final concentration of 10 µM and imaged using nominal 450/485 nm excitation/emission settings. IPG-4 AM was used at a final concentration of 3 µM. These measurements were interpreted as relative fluorescence dynamics, not as calibrated membrane voltage or absolute potassium concentration.

### 4.8. Lempel-Ziv pattern complexity and O-information analysis

#### Image pre-processing

Images of individual wells were first extracted from the wide-field microscopy images using the Cellpose segmentation algorithm [59], after which the green channel, corresponding to the fluorescence signal, was isolated. The images were then binarized at the median pixel value, producing an approximately maximum-entropy signal with similar numbers of 0- and 1-valued pixels. Following previous work in signal processing [60, 61], the two-dimensional binary data were flattened into a one-dimensional binary string.

#### Computing the Lempel-Ziv complexity

Lempel-Ziv complexity (LZC) [62] relates the complexity of a discrete string of symbols to the size of its compressed representation. Intuitively, computing the LZC of a string *X*involves iterating through the array element by element and constructing a dictionary of substrings of *X*, with a new entry added whenever a previously unobserved substring is encountered. The size of the resulting dictionary is proportional to the number of distinct patterns represented in *X*.

Since individual wells contained different numbers of pixels, LZC values were normalized to a 0-1 interval using a maximum-entropy null model. Following previous work [60, 61], the LZC of each string was recalculated after randomly shuffling its elements to disrupt cross-pixel correlations. This procedure was repeated five times, and the observed LZC was normalized by the expected value of the null LZCs. Increased complexity in this context does not necessarily imply increased visual interestingness, because white noise has maximal LZC.

LZC was computed using the implementation provided in the Syntropy Python package (https://github.com/thosvarley/syntropy).

#### Higher-order interaction analysis

To explore interactions among embryos or Xenobots, bacteria, and their shared environment, we computed the O-information [63] among normalized LZC, treated as a continuous variable; KCl concentration, treated as a continuous variable; and the binary presence or absence of an embryo or Xenobot, treated as a discrete random variable. For a three-variable system, the O-information can be expressed as a linear combination of mutual informations:

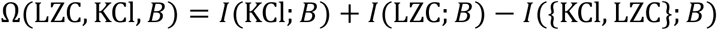

where mutual information is defined as:

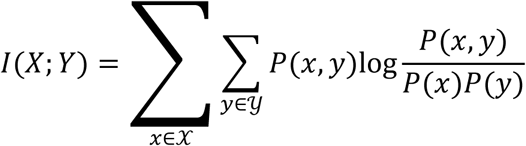

Since normalized LZC was not normally distributed, 1 − LZCwas log-transformed to make the variable more amenable to Gaussian estimators.

A negative O-information value reflects irreducible synergistic information about entity presence or absence that is contained in the joint state of KCl concentration and LZC but is inaccessible when either predictor is considered independently.

Mixed discrete-continuous mutual information was estimated using the identity:

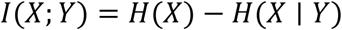

Assuming that *X*was sampled from a multivariate Gaussian distribution, joint and conditional entropies were estimated from the covariance matrices of the complete dataset and from the covariance matrices obtained after conditioning on each state *Y* = *y*in the support of *Y*.

The statistical significance of the O-information was assessed using a permutation null distribution with 25,000 replications, in which all variables were randomized independently.

### 4.9. Machine learning analysis of distributed bacterial patterns

We used supervised learning to determine whether bacterial spatial patterns were predictive of the presence and identity of a *Xenopus*-derived entity. The model was trained on time-lapse fluorescence microscopy videos of bacterial cultures containing a Xenobot, a *Xenopus* embryo, or no biological entity.

To prevent the classifier from relying directly on the morphology of embryos or Xenobots, entity silhouettes were detected automatically and excluded from the input data. Individual wells were first identified in each frame by thresholding in HSV color space and segmenting the resulting green connected components. Within each well, the largest contour in the binarized image was taken to represent the embryo or Xenobot silhouette. Image patches overlapping this contour were removed from the dataset. Manual inspection confirmed that this procedure accurately detected the biological entities in all analyzed videos.

Videos from six wells for each entity type and KCl concentration were assigned to the training dataset, and videos from an additional six wells for each entity type and KCl concentration were reserved as a held-out test dataset. Each frame was divided into a 5x5 grid, producing 25 image patches. After removal of patches overlapping embryos or Xenobots, the dataset was balanced by randomly subsampling patches so that each class contributed an equal number of examples. The resulting dataset contained 427,500 patches. Videos, not individual patches, were divided between the training and held-out datasets, ensuring that patches originating from the same video were not represented in both sets. Consequently, successful classification required the model to learn bacterial-pattern features that generalized to previously unseen videos.

A convolutional neural network was trained to classify each image patch into one of three categories: Xenobot, embryo, or no biological entity. The network contained three convolutional layers, each followed by batch normalization, max pooling, and a rectified linear unit activation. The convolutional feature extractor was followed by a classifier containing two fully connected layers. The model was optimized using cross-entropy loss and the Adam optimizer with a learning rate of 0.001. Training was performed for 30 epochs, by which point model performance had reached a stable plateau.

### 4.10. Computational and statistical analysis

Data are presented as mean±SEM unless otherwise stated. Statistical tests were applied to independent well-, sample-, experimental-repeat-, or data-split-level values, as appropriate. Tests were two-sided except for the two directional comparisons in Figure 5 specified in the corresponding Results section. The statistical model, test, analysis window, and multiple-comparison correction used for each experiment are reported in the corresponding figure or Results section. Statistical significance was defined as *p<0.05* after the stated correction.

Image processing, quantitative image analysis, and statistical analyses for Figures 1-7 were performed in Python 3.13.5 using NumPy 2.3.5, pandas 2.2.3, SciPy 1.17.0, statsmodels 0.14.6, scikit-image 0.26.0, and opencv-python-headless 4.13.0.92. Quantitative plots and analysis workbooks were generated using Matplotlib 3.10.8 and openpyxl 3.1.5. Lempel-Ziv complexity and O-information analyses for Figure 8 and machine-learning analyses for Figure 9 were performed using separate Python pipelines, as described in Sections 4.8 and 4.9. Analysis-specific procedures for randomization, permutation testing, class balancing, and held-out evaluation are detailed in those sections.

**Figure 8.**
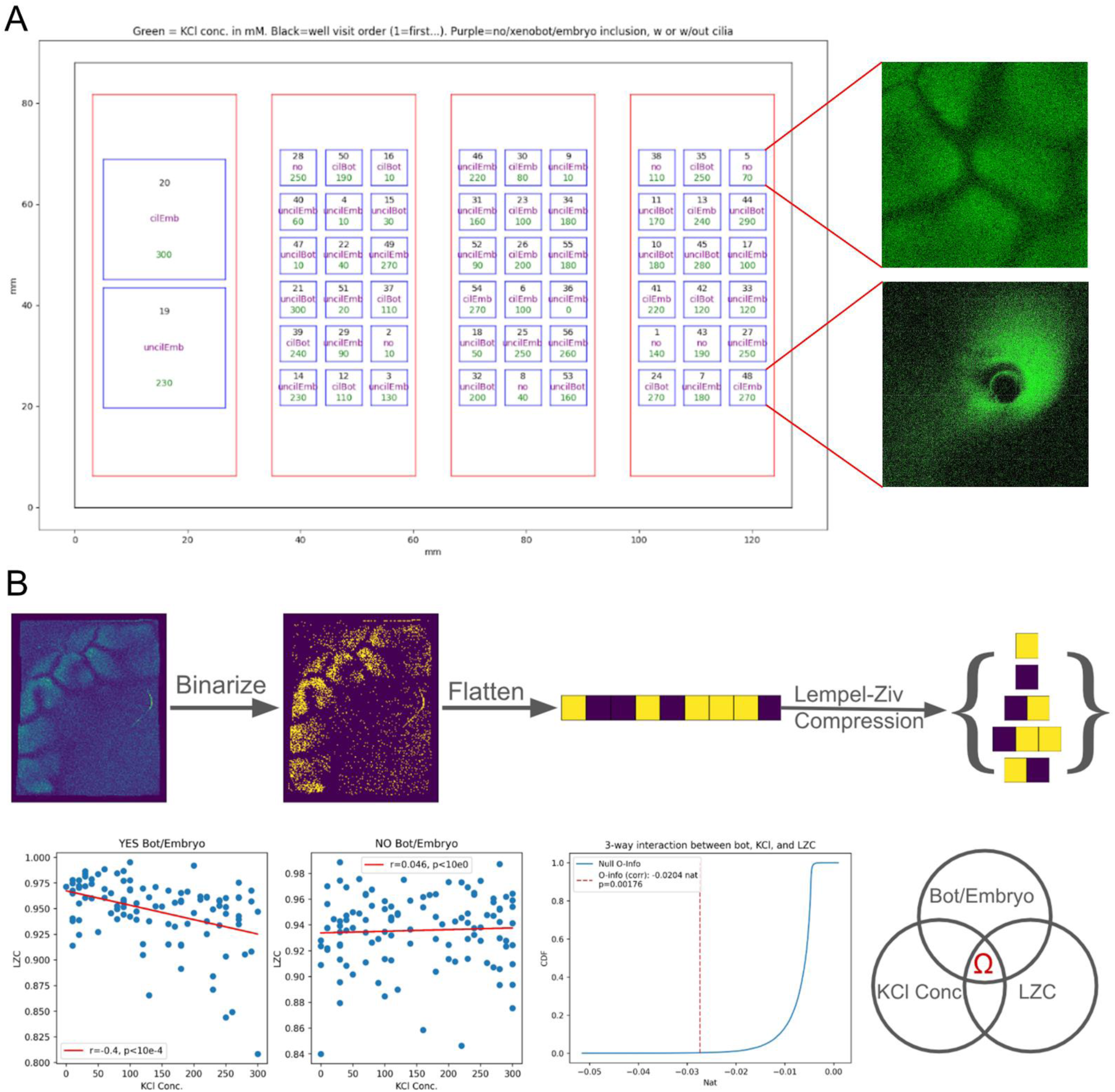
Xenobot/embryo presence modulates the relationship between ionic stress and bacterial pattern complexity. A. AI-generated schematic of the experimental design showing the arrangement of four microscope slides containing wells of different sizes on the motorized stage. Green numbers indicate KCl concentration (mM), black numbers indicate well visit order during imaging (1=first), and purple labels indicate sample identity: bacteria alone (“no”) or bacteria with Xenobot/embryo constructs, with or without cilia. Representative fluorescence images of bacterial patterns from selected wells are shown at right. B. Top row: Workflow for normalized Lempel-Ziv complexity (LZC) analysis of bacterial pattern images. Raw fluorescence images were binarized using a threshold of approximately twice the median image intensity, converted into a one-dimensional binary sequence by row-wise flattening, and compressed using the classical Lempel-Ziv algorithm. Complexity was then normalized relative to the expected compression of a maximum-entropy binary string of the same length. Bottom row: Analysis of the interaction between Xenobot/embryo presence, KCl concentration, and bacterial pattern complexity. In samples containing a Xenobot or embryo, increasing KCl was associated with a significant decrease in normalized LZC (r=-0.40, p<10^-4^), whereas in samples lacking a Xenobot/embryo, no meaningful relationship was observed (r=0.046, n.s.). The three-way dependency among Xenobot/embryo status, KCl concentration, and LZC was quantified using the O-information [63], yielding a negative value (Ω=-0.0204 nat, p=0.00176), consistent with a synergistic interaction. These results indicate that the effect of ionic stress on bacterial collective organization emerges specifically in the presence of the living multicellular construct.

**Figure 9.**
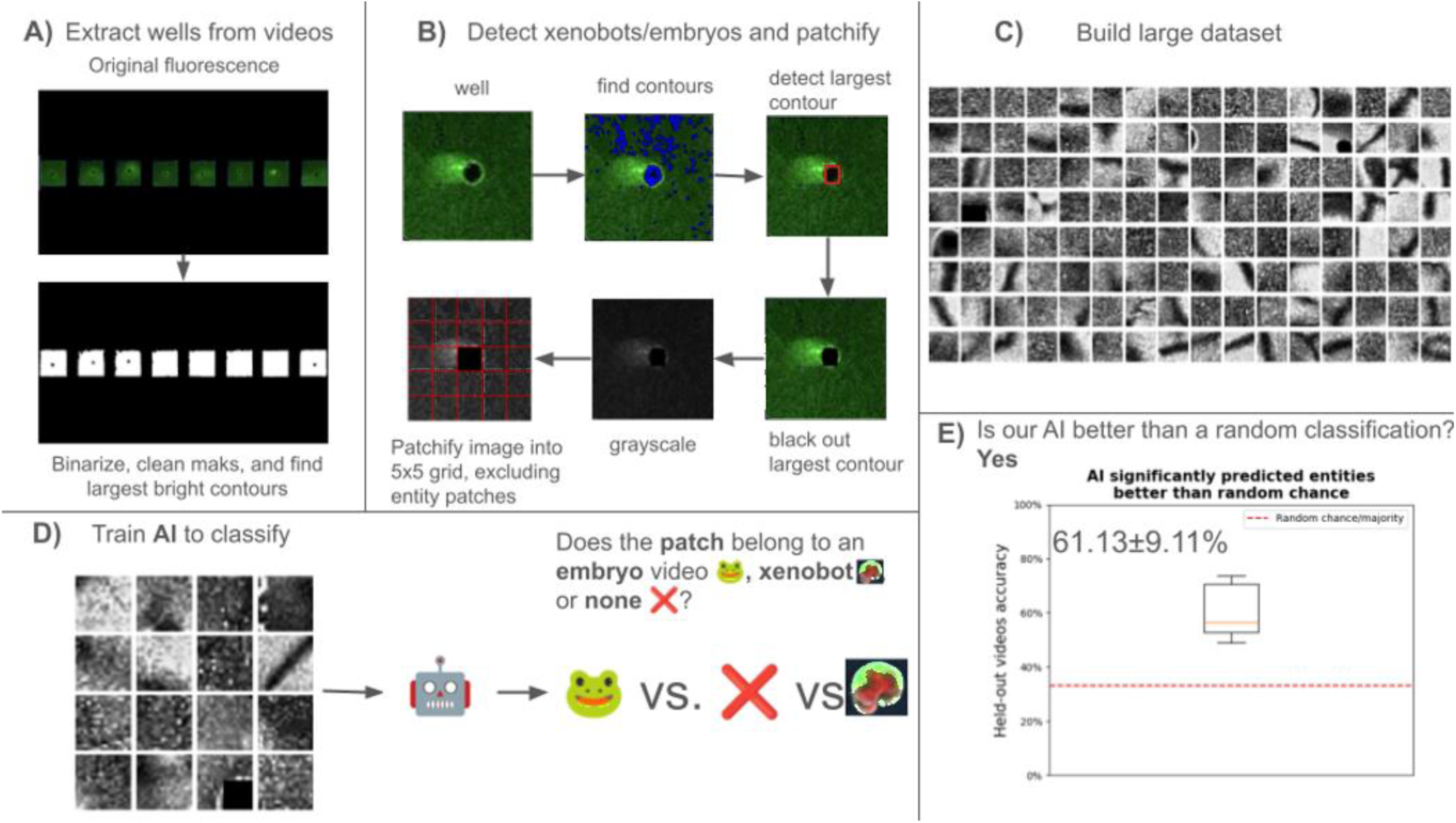
AI detects distributed bacterial pattern signatures of hidden *Xenopus* embryo or Xenobot presence. A. Individual wells were detected from fluorescence frames by binarization, cleaning of the white masks, and identifying the disconnected largest contours (one per well). B. For each well, *Xenopus* embryos and Xenobots were detected by identifying the black contours (highlighted in blue in the second frame), retaining the largest one, and blacking out its bounding rectangle. For more efficient neural network processing, the frames were then grayscaled and divided into patches. To prevent the supervised model from using direct visual information from embryos or Xenobots, the patches overlapping with the detected embryo/Xenobot rectangles were excluded from the dataset. In this way, the model was forced to rely on bacterial spatial organization instead of constructing silhouettes, position, or residual morphology. C. Patches from videos containing Xenobots, embryos, or bacteria only were assembled into a large training dataset across KCl conditions. D. The model was trained to classify whether bacterial-pattern patches originated from Xenobot-containing, embryo-containing, or bacteria-only cultures and tested on held-out videos. E. Despite removal of direct embryo/Xenobot visual information, classification accuracy on held-out videos was significantly above the 3-class majority-classifier baseline of 33%, reaching 61.13±9.11% accuracy across train/test splits and random seeds (Mann-Whitney, p<0.001). These results indicate that bacterial spatial organization contains distributed signatures of *Xenopus*-derived constructs even when the constructs themselves are hidden from view.

## Acknowledgments

The authors thank Patrick McMillen, Doug Blackiston, Vaibhav Pai, and many members of the Levin Lab for helpful discussions, advice, and technical guidance; Erin Switzer for animal husbandry; Rakela Colon for laboratory management and support; Ann Amanda Karim for project management and coordination; and Tomika Gotch for assistance with manuscript preparation.

## References

1. Hughes, D.T. and V. Sperandio, Inter-kingdom signalling: communication between bacteria and their hosts. Nature Reviews Microbiology, 2008. 6(2): p. 111–120.

2. Pacheco, A.R. and V. Sperandio, Inter-kingdom signaling: chemical language between bacteria and host. Current Opinion in Microbiology, 2009. 12(2): p. 192–198.

3. McFall-Ngai, M., et al., Animals in a bacterial world, a new imperative for the life sciences. Proceedings of the National Academy of Sciences of the United States of America, 2013. 110(9): p. 3229–3236.

4. McMillen, P. and M. Levin, Collective intelligence: A unifying concept for integrating biology across scales and substrates. Communications Biology, 2024. 7(1): p. 378.

5. Heylighen, F., Self-organization in communicating groups: The emergence of coordination, shared references and collective intelligence. In: Massip-Bonet, À. and A. Bastardas-Boada, eds. Complexity Perspectives on Language, Communication and Society. Springer, 2013: p. 117–149.

6. Couzin, I., Collective minds. Nature, 2007. 445(7129): p. 715.

7. Strassmann, J.E. and D.C. Queller, The social organism: congresses, parties, and committees. Evolution, 2010. 64(3): p. 605–616.

8. Ben-Jacob, E., Y. Aharonov, and Y. Shapira, Bacteria harnessing complexity. Biofilms, 2004. 1(4): p. 239–263.

9. Ben-Jacob, E., Learning from bacteria about natural information processing. Annals of the New York Academy of Sciences, 2009. 1178: p. 78–90.

10. Ben-Jacob, E., Y. Shapira, and A.I. Tauber, Seeking the foundations of cognition in bacteria: From Schrödinger’s negative entropy to latent information. Physica A: Statistical Mechanics and Its Applications, 2006. 359: p. 495–524.

11. Hellingwerf, K.J., Bacterial observations: a rudimentary form of intelligence? Trends in Microbiology, 2005. 13(4): p. 152–158.

12. Lyon, P., The cognitive cell: bacterial behavior reconsidered. Frontiers in Microbiology, 2015. 6: p. 264.

13. Shapiro, J.A., The significances of bacterial colony patterns. BioEssays, 1995. 17(7): p. 597–607.

14. Ben-Jacob, E., I. Cohen, and D.L. Gutnick, Cooperative organization of bacterial colonies: from genotype to morphotype. Annual Review of Microbiology, 1998. 52: p. 779–806.

15. O’Toole, G., H.B. Kaplan, and R. Kolter, Biofilm formation as microbial development. Annual Review of Microbiology, 2000. 54: p. 49–79.

16. Branda, S.S., et al., Genes involved in formation of structured multicellular communities by Bacillus subtilis. Journal of Bacteriology, 2004. 186(12): p. 3970–3979.

17. Kearns, D.B., et al., A master regulator for biofilm formation by Bacillus subtilis. Molecular Microbiology, 2005. 55(3): p. 739–749.

18. Branda, S.S., et al., A major protein component of the Bacillus subtilis biofilm matrix. Molecular Microbiology, 2006. 59(4): p. 1229–1238.

19. Romero, D., et al., Amyloid fibers provide structural integrity to Bacillus subtilis biofilms. Proceedings of the National Academy of Sciences of the United States of America, 2010. 107(5): p. 2230–2234.

20. Vlamakis, H., et al., Sticking together: building a biofilm the Bacillus subtilis way. Nature Reviews Microbiology, 2013. 11(3): p. 157–168.

21. Prindle, A., et al., Ion channels enable electrical communication in bacterial communities. Nature, 2015. 527(7576): p. 59–63.

22. Humphries, J., et al., Species-independent attraction to biofilms through electrical signaling. Cell, 2017. 168(1-2): p. 200–209.e12.

23. Liu, J., et al., Coupling between distant biofilms and emergence of nutrient time-sharing. Science, 2017. 356(6338): p. 638–642.

24. Larkin, J.W., et al., Signal percolation within a bacterial community. Cell Systems, 2018. 7(2): p. 137–145.e3.

25. Martinez-Corral, R., et al., Bistable emergence of oscillations in growing Bacillus subtilis biofilms. Proceedings of the National Academy of Sciences of the United States of America, 2018. 115(36): p. E8333–E8340.

26. Chou, K., et al., A segmentation clock patterns cellular differentiation in a bacterial biofilm. Cell, 2022. 185(1): p. 145–157.e13.

27. Nyholm, S.V., et al., Roles of Vibrio fischeri and nonsymbiotic bacteria in the dynamics of mucus secretion during symbiont colonization of the Euprymna scolopes light organ. Applied and Environmental Microbiology, 2002. 68(10): p. 5113–5122.

28. Nyholm, S.V. and M.J. McFall-Ngai, Dominance of Vibrio fischeri in secreted mucus outside the light organ of Euprymna scolopes: the first site of symbiont specificity. Applied and Environmental Microbiology, 2003. 69(7): p. 3932–3937.

29. Jemielita, M., et al., Spatial and temporal features of the growth of a bacterial species colonizing the zebrafish gut. mBio, 2014. 5(6): p. e01751–14.

30. Earle, K.A., et al., Quantitative imaging of gut microbiota spatial organization. Cell Host & Microbe, 2015. 18(4): p. 478–488.

31. Wheeler, K.M., et al., Mucin glycans attenuate the virulence of Pseudomonas aeruginosa in infection. Nature Microbiology, 2019. 4(12): p. 2146–2154.

32. Wiles, T.J., et al., Swimming motility of a gut bacterial symbiont promotes resistance to intestinal expulsion and enhances inflammation. PLOS Biology, 2020. 18(3): p. e3000661.

33. Smith, T.J., et al., A mucin-regulated adhesin determines the spatial organization and inflammatory character of a bacterial symbiont in the vertebrate gut. Cell Host & Microbe, 2023. 31(8): p. 1371–1385.e6.

34. Rossy, T., et al., Pseudomonas aeruginosa type IV pili actively induce mucus contraction to form biofilms in tissue-engineered human airways. PLOS Biology, 2023. 21(8): p. e3002209.

35. Beauregard, P.B., et al., Bacillus subtilis biofilm induction by plant polysaccharides. Proceedings of the National Academy of Sciences of the United States of America, 2013. 110(17): p. E1621–E1630.

36. Allard-Massicotte, R., et al., Bacillus subtilis early colonization of Arabidopsis thaliana roots involves multiple chemotaxis receptors. mBio, 2016. 7(6): p. e01664–16.

37. Bais, H.P., R. Fall, and J.M. Vivanco, Biocontrol of Bacillus subtilis against infection of Arabidopsis roots by Pseudomonas syringae is facilitated by biofilm formation and surfactin production. Plant Physiology, 2004. 134(1): p. 307–319.

38. Rapsinski, G.J., et al., Pseudomonas aeruginosa senses and responds to epithelial potassium flux via Kdp operon to promote biofilm. PLOS Pathogens, 2024. 20(5): p. e1011453.

39. Alegado, R.A., et al., A bacterial sulfonolipid triggers multicellular development in the closest living relatives of animals. eLife, 2012. 1: p. e00013.

40. Sundelacruz, S., M. Levin, and D.L. Kaplan, Role of membrane potential in the regulation of cell proliferation and differentiation. Stem Cell Reviews and Reports, 2009. 5(3): p. 231–246.

41. Levin, M., Molecular bioelectricity: how endogenous voltage potentials control cell behavior and instruct pattern regulation in vivo. Molecular Biology of the Cell, 2014. 25(24): p. 3835–3850.

42. Bates, E., Ion channels in development and cancer. Annual Review of Cell and Developmental Biology, 2015. 31: p. 231–247.

43. Levin, M., G. Pezzulo, and J.M. Finkelstein, Endogenous bioelectric signaling networks: Exploiting voltage gradients for control of growth and form. Annual Review of Biomedical Engineering, 2017. 19: p. 353–387.

44. Levin, M., Bioelectric signaling: Reprogrammable circuits underlying embryogenesis, regeneration, and cancer. Cell, 2021. 184(8): p. 1971–1989.

45. Harris, M.P., Bioelectric signaling as a unique regulator of development and regeneration. Development, 2021. 148(10): p. dev180794.

46. Adams, D.S., A. Masi, and M. Levin, H+ pump-dependent changes in membrane voltage are an early mechanism necessary and sufficient to induce *Xenopus* tail regeneration. Development, 2007. 134(7): p. 1323–1335.

47. Tseng, A., et al., Induction of vertebrate regeneration by a transient sodium current. Journal of Neuroscience, 2010. 30(39): p. 13192–13200.

48. Pai, V.P., et al., Transmembrane voltage potential controls embryonic eye patterning in *Xenopus* laevis. Development, 2012. 139(2): p. 313–323.

49. Chernet, B.T. and M. Levin, Transmembrane voltage potential is an essential cellular parameter for the detection and control of tumor development in a *Xenopus* model. Disease Models & Mechanisms, 2013. 6(3): p. 595–607.

50. Levin, M., Endogenous bioelectrical networks store non-genetic patterning information during development and regeneration. Journal of Physiology, 2014. 592(11): p. 2295–2305.

51. Levin, M., The computational boundary of a “self”: Developmental bioelectricity drives multicellularity and scale-free cognition. Frontiers in Psychology, 2019. 10: p. 2688.

52. Kriegman, S., et al., A scalable pipeline for designing reconfigurable organisms. Proceedings of the National Academy of Sciences of the United States of America, 2020. 117(4): p. 1853–1859.

53. Blackiston, D., et al., A cellular platform for the development of synthetic living machines. Science Robotics, 2021. 6(52): p. eabf1571.

54. Kriegman, S., et al., Kinematic self-replication in reconfigurable organisms. Proceedings of the National Academy of Sciences of the United States of America, 2021. 118(49): p. e2112672118.

55. Pai, V.P., et al., Basal Xenobot transcriptomics reveals changes and novel control modality in cells freed from organismal influence. Communications Biology, 2025. 8(1): p. 646.

56. Nieuwkoop, P.D. and J. Faber, eds., Normal Table of Xenopus laevis (Daudin): A Systematical and Chronological Survey of the Development from the Fertilized Egg till the End of Metamorphosis. 2nd ed. Amsterdam: North-Holland Publishing Company, 1967.

57. Irnov, I. and W.C. Winkler, A regulatory RNA required for antitermination of biofilm and capsular polysaccharide operons in Bacillales. Molecular microbiology, 2010. 76(3): p. 559–575.

58. Liu, J., et al., Metabolic co-dependence gives rise to collective oscillations within biofilms. Nature, 2015. 523(7562): p. 550–554.

59. Stringer, C., et al., Cellpose: a generalist algorithm for cellular segmentation. Nature Methods, 2021. 18(1): p. 100–106.

60. Schartner, M., et al., Complexity of multi-dimensional spontaneous EEG decreases during propofol induced general anaesthesia. PLOS ONE, 2015. 10(8): p. e0133532.

61. Schartner, M.M., et al., Increased spontaneous MEG signal diversity for psychoactive doses of ketamine, LSD and psilocybin. Scientific Reports, 2017. 7: p. 46421.

62. Lempel, A. and J. Ziv, On the complexity of finite sequences. IEEE Transactions on Information Theory, 1976. 22(1): p. 75–81.

63. Rosas, F.E., et al., Quantifying high-order interdependencies via multivariate extensions of the mutual information. Physical Review E, 2019. 100(3-1): p. 032305.

64. Holtmann, G., et al., KtrAB and KtrCD: two K+ uptake systems in Bacillus subtilis and their role in adaptation to hypertonicity. Journal of Bacteriology, 2003. 185(4): p. 1289–1298.

65. Gundlach, J., et al., Control of potassium homeostasis is an essential function of the second messenger cyclic di-AMP in Bacillus subtilis. Science Signaling, 2017. 10(475): p. eaal3011.

66. Varley, T.F., A synergistic perspective on multivariate computation and causality in complex systems. Entropy, 2024. 26(10): p. 883.

67. Dixon, T.A., T.C. Williams, and I.S. Pretorius, Sensing the future of bio-informational engineering. Nature Communications, 2021. 12(1): p. 388.

68. Levin, M., Darwin’s agential materials: evolutionary implications of multiscale competency in developmental biology. Cellular and Molecular Life Sciences, 2023. 80(6): p. 142.

69. Pfeifer, R., M. Lungarella, and F. Iida, Self-organization, embodiment, and biologically inspired robotics. Science, 2007. 318(5853): p. 1088–1093.

70. Bongard, J., Morphological change in machines accelerates the evolution of robust behavior. Proceedings of the National Academy of Sciences of the United States of America, 2011. 108(4): p. 1234–1239.

71. Cheney, N., et al., Scalable co-optimization of morphology and control in embodied machines. Journal of the Royal Society Interface, 2018. 15(143): p. 20170937.

72. Müller, V.C. and M. Hoffmann, What is morphological computation? On how the body contributes to cognition and control. Artificial Life, 2017. 23(1): p. 1–24.

73. Chandrasekaran, S.N., et al., Image-based profiling for drug discovery: due for a machine-learning upgrade? Nature Reviews Drug Discovery, 2021. 20(2): p. 145–159.

74. Caicedo, J.C., et al., Data-analysis strategies for image-based cell profiling. Nature Methods, 2017. 14(9): p. 849–863.

75. Lu, H.Y., et al., Multi-scale neural decoding and analysis. Journal of Neural Engineering, 2021. 18(4): p. 045013.

76. Huth, A.G., et al., Decoding the semantic content of natural movies from human brain activity. Frontiers in Systems Neuroscience, 2016. 10: p. 81.

77. Pio-Lopez, L., G. Pezzulo, and M. Levin, Scale-free niche construction: expanding agent- microenvironment co-development to unconventional substrates. OSF Preprints, 2025. doi: 10.31219/osf.io/ytg35_v1.

78. Laland, K.N., et al., The extended evolutionary synthesis: its structure, assumptions and predictions. Proceedings of the Royal Society B: Biological Sciences, 2015. 282(1813): p. 20151019.

79. Chiu, L. and S.F. Gilbert, The birth of the holobiont: Multi-species birthing through mutual scaffolding and niche construction. Biosemiotics, 2015. 8(2): p. 191–210.

