## Supplementary Figures for "Living multicellular systems induce decodable spatial patterns in bacterial collectives"

### Supplements

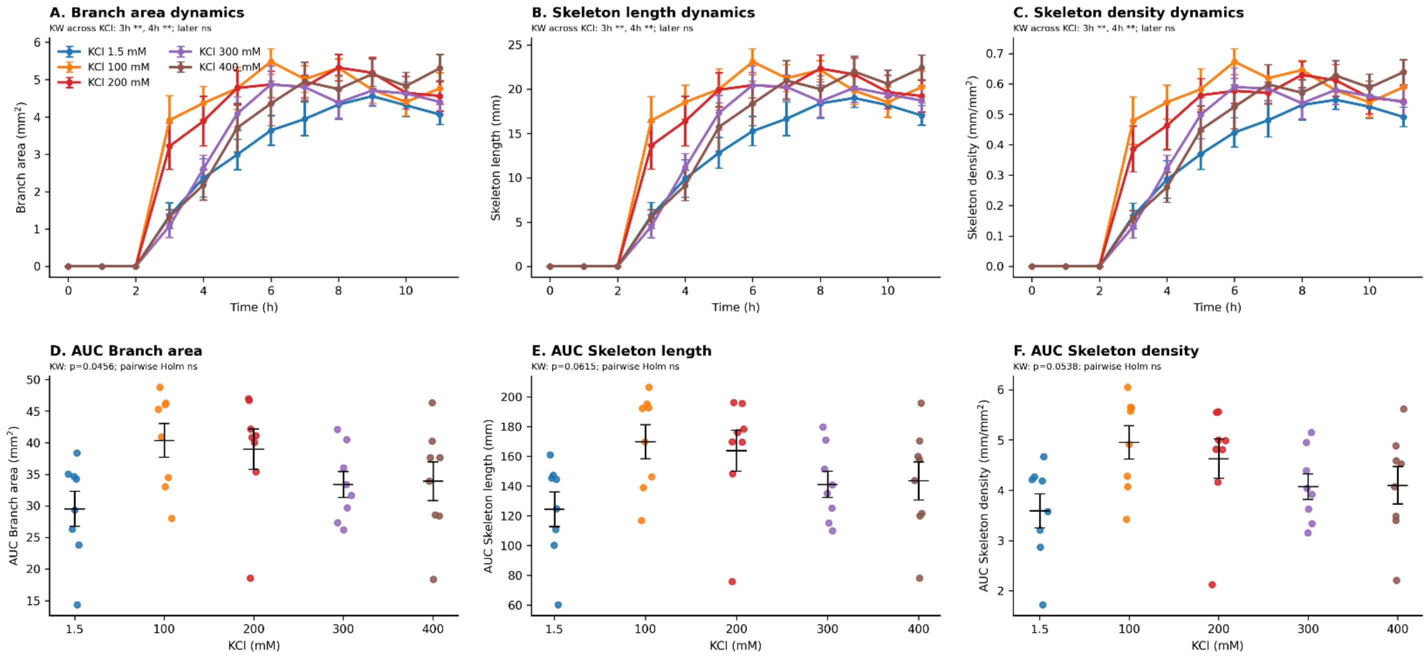


Supplement Figure 1. Potassium concentration does not alter branching dynamics in target-free wells.

A-C. Branching-network dynamics in target-free wells across KCl concentrations from 1.5 to 400 mM. Branch area (A), branch skeleton length (B), and skeleton density (C) increased over time in all conditions, indicating progressive branching-network formation in the absence of a target. Early time-point differences were detected across KCl concentrations, but later time points did not show a consistent concentration-dependent separation.

D-F. Area-under-the-curve (AUC) analysis of branch area (D), skeleton length (E), and skeleton density (F) across the full 0-11 h time course. AUC was calculated for each well by trapezoidal integration of hourly measurements. Branch area AUC showed a borderline overall KCl effect (Kruskal-Wallis p=0.0456), whereas skeleton length AUC and skeleton density AUC did not reach significance (p=0.0615 and p=0.0538, respectively). All pairwise comparisons between KCl concentrations were non-significant after Holm correction. Data are mean±SEM for time-course plots and individual wells with mean±SEM for AUC plots; n=8 wells per condition. This target-free control showes that potassium alone does not robustly suppress branching-network formation.

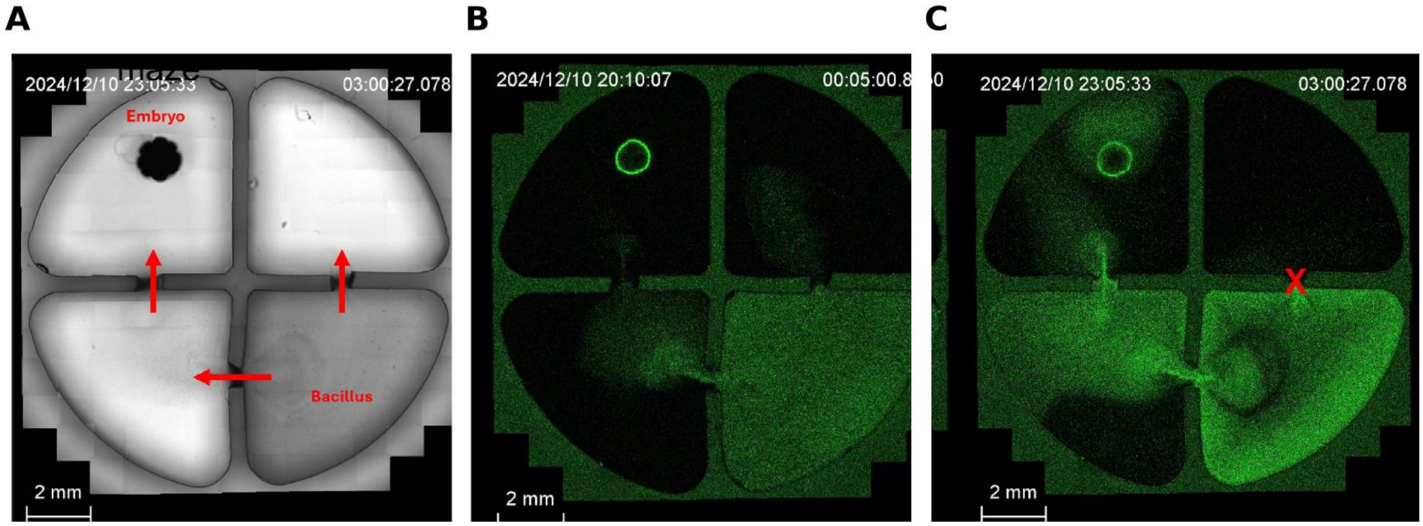


Supplement Figure 2. Exploratory maze-like assay of embryo-associated bacterial attraction in a structured environment.

A. Bright-field overview of the multi-chamber maze geometry. An embryo was placed in the upper-left chamber, while *B. subtilis* $P_{hyp}$-*yfp* bacteria were introduced into the bottom-right chamber. Red arrows indicate possible routes through the chamber connections.

B. Early YFP fluorescence image showing initial bacterial distribution shortly after introduction into the maze. A small bacterial population transiently entered the upper-right empty chamber, while bacterial signal was also present along the route toward the embryo-containing chamber.

C. Later YFP fluorescence image from the same assay showing preferential bacterial accumulation toward the embryo-containing chamber and reduced signal in the upper-right empty chamber. The red "X" marks the empty chamber region from which bacterial signal was reduced over time. Scale bars, 2 mm.

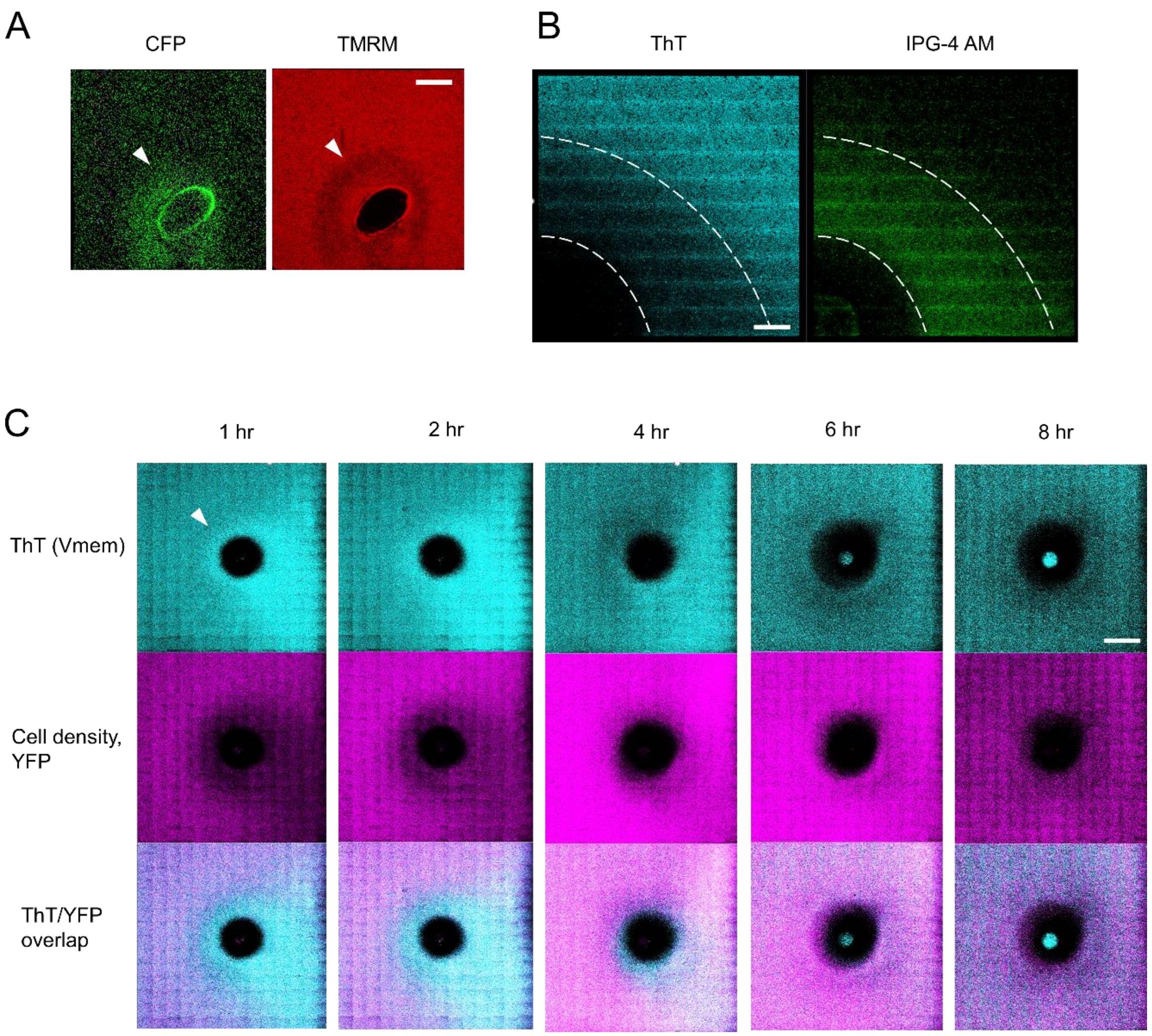


Supplement Figure 3. Embryo proximity is associated with bacterial bioelectrical and potassium dynamics.

A. Bacterial attraction transiently coincides with membrane-potential-associated depolarization. Fluorescence images show motile *B. subtilis* $P_{hyp}$-*cfp* cells forming an attraction halo around a *Xenopus* embryo in high KCl medium (300 mM; left). TMRM imaging of the same field shows a transient depolarization-associated signal near the attraction region (right). White arrows mark bacterial accumulation and the corresponding TMRM signal. *Xenopus* embryo, NF~22. Scale bar, 1 mm.

B. Membrane-potential-associated Thioflavin signal and intracellular-accessible potassium signal near the embryo in non-motile bacteria. *Δhag* $P_{hyp}$-*yfp* cells were used to minimize bacterial accumulation around the embryo, allowing local signal changes to be examined independently of attraction-halo formation. Thioflavin reports relative membrane-potential-associated changes, whereas IPG-4 AM reports intracellular-accessible potassium-associated signal. KCl, 0.2 mM. Scale bar, 100 µm.

C. Temporal relationship between membrane-potential-associated Thioflavin signal and bacterial cell density in non-motile bacteria near the embryo. Time-lapse snapshots show Thioflavin signal (top row), bacterial density visualized by YFP reporter fluorescence (middle row), and spatial overlap between the two channels (bottom row) at 1, 2, 4, 6, and 8 hr. Because *Δhag* $P_{hyp}$-*yfp* cells do not form motility-dependent attraction halos, this assay separates embryo-associated bioelectrical changes from bacterial accumulation. The white arrow marks a region of hyperpolarization-associated signal near the embryo. KCl 0.2 mM  Scale bar, 1 mm.

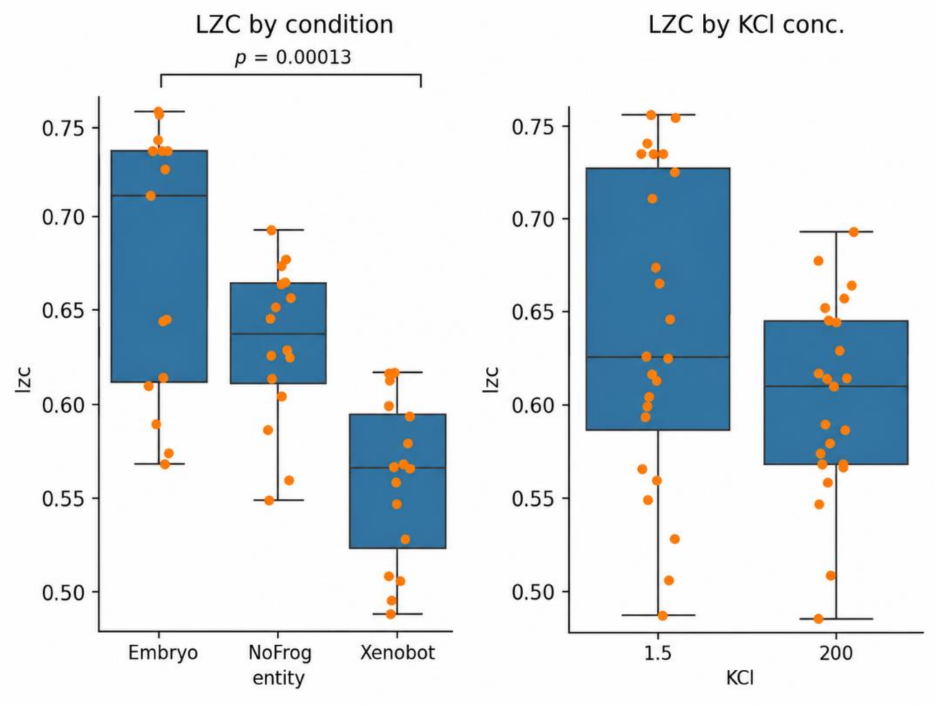


Supplement Figure 4. Pattern complexity differs across entity conditions in the secondary dataset. Normalized Lempel-Ziv complexity (LZC) of bacterial spatial patterns in the secondary dataset, collected using fixed well sizes and two KCl concentrations (1.5 mM and 200 mM). Left: LZC grouped by biological condition: embryo, no-frog control, and Xenobot. A Kruskal-Wallis test detected a significant overall difference among conditions ($H = 20.58$, $p < 10^{-4}$). Post hoc pairwise comparison using the Mann-Whitney U test showed a significant difference between the embryo and Xenobot conditions ($U = 23$, $p = 0.00013$), with embryo-associated patterns exhibiting higher normalized LZC than Xenobot-associated patterns. Right: The same LZC values grouped by KCl concentration (1.5 mM vs. 200 mM), shown to illustrate the distribution of pattern complexity across the two ionic conditions in this dataset. Boxplots show median and interquartile range; whiskers show the data range excluding outliers, and overlaid points represent individual samples.

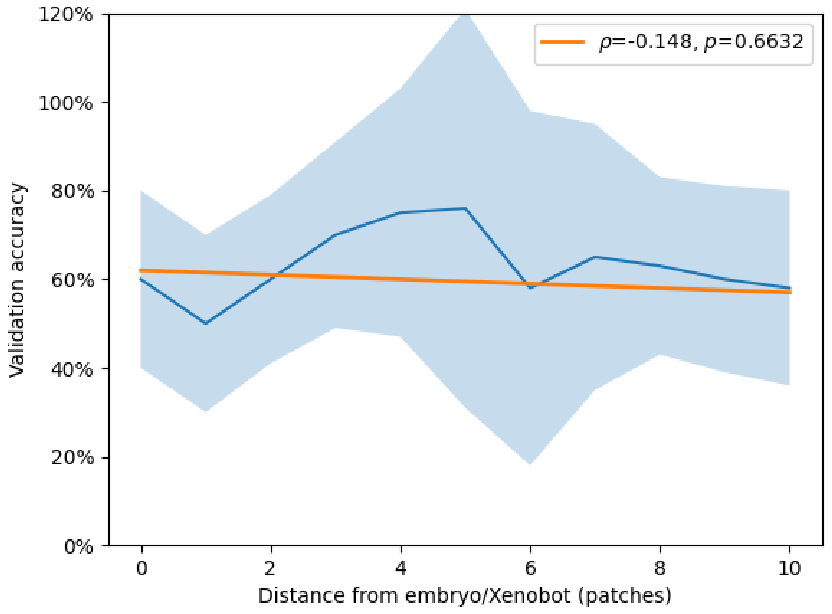

Supplement Figure 5. The classification accuracy of the supervised AI is not affected by distance from the *Xenopus* embryo or Xenobot.

Accuracy on held-out patches, measured as mean±std across splits and random seeds. The orange line is the linear regression fit, with Pearson's coefficient and its p-value in the legend. Since there's no significant relationship, the AI learned spatially distributed bacterial patterns that differentiated between biological entities.
